# Morphogenesis in *Trypanosoma cruzi* epimastigotes proceeds via a highly asymmetric cell division

**DOI:** 10.1101/2023.05.24.542100

**Authors:** Paul C. Campbell, Christopher L. de Graffenried

**Author notes:** Address correspondence to Christopher de Graffenried.

## Abstract

*Trypanosoma cruzi* is a protist parasite that is the causative agent of Chagas’ disease, a neglected tropical disease endemic to the Americas. *T. cruzi* cells are highly polarized and undergo morphological changes as they cycle within their insect and mammalian hosts. Work on related trypanosomatids has described cell division mechanisms in several life-cycle stages and identified a set of essential morphogenic proteins that serve as markers for key events during trypanosomatid division. Here, we use Cas9-based tagging of morphogenic genes, live-cell imaging, and expansion microscopy to study the cell division mechanism of the insect-resident epimastigote form of *T. cruzi,* which represents an understudied trypanosomatid morphotype. We find that *T. cruzi* epimastigote cell division is highly asymmetric, producing one daughter cell that is significantly smaller than the other. Daughter cell division rates differ by 4.9 h, which may be a consequence of this size disparity. Many of the morphogenic proteins identified in *T. brucei* have altered localization patterns in *T. cruzi* epimastigoes, which may reflect fundamental differences in the cell division mechanism of this life cycle stage, which widens and shortens the cell body to accommodate the duplicated organelles and cleavage furrow rather than elongating the cell body along the long axis of the cell, as is the case in life-cycle stages that have been studied in *T. brucei*. This work provides a foundation for further investigations of *T. cruzi* cell division and shows that subtle differences in trypansomatid cell morphology can alter how these parasites divide.

**Author Summary:** *Trypanosoma cruzi* causes Chagas’ disease, which is among the most neglected of tropical diseases, affecting millions of people in South and Central America along with immigrant populations around the world. *T. cruzi* is related to other important pathogens such as *Trypanosoma brucei* and *Leishmania spp,* which have been the subject of molecular and cellular characterizations that have provided an understanding of how these organisms shape their cells and undergo division. Work in *T. cruzi* has lagged due to an absence of molecular tools for manipulating the parasite and the complexity of the original published genome; these issues have recently been resolved. Building on work in *T. brucei*, we have studied the localization of key cell cycle proteins and quantified changes in cell shape during division in an insect-resident form of *T. cruzi*. This work has uncovered unique adaptations to the cell division process in *T. cruzi* and provides insight into the range of mechanisms this family of important pathogens can employ to colonize their hosts.

## Introduction

The Trypanosomatidae family encompasses parasitic species that inhabit a broad range of hosts including plants, insects, fish, and mammals[1–3]. These parasites tune core pathways including their metabolism and cell surface proteomes to survive within varied host environments[4–6]. Trypanosomatids have also adapted their cellular morphology to facilitate processes such as evasion of the host immune system, attachment to epithelial layers, and proliferation within host cells[7,8]. These morphological states have required adaptations to other essential cellular pathways such as cell division and endocytosis [9,10]. In recent years, work on the parasites *Trypanosoma brucei* and *Leishmania spp* has uncovered a range of trypanosomatid cell division mechanisms and identified key regulators, which include conserved eukaryotic proteins and trypanosomatid-specific components[10,11]. *Trypanosoma cruzi*, the causative agent of Chagas’ disease, is a significant human health burden, with over 7 million chronic infections primarily in Latin America and 50,000 deaths annually[12]. The cellular biology of *T. cruzi* has not been studied as extensively as other trypanosomatids, primarily due to the complexity of the original sequenced genome and the lack of tools such as RNAi to probe essential gene function[13,14]. Recent genome sequencing of several *T. cruzi* strains and the development of Cas9 editing systems have remedied many of these problems [14–17].

Trypanosomatids share a set of organelles that facilitate infection and subsequent survival within their hosts[18]. As part of the Kinetoplastea class, trypanosomatids have a mitochondrial DNA aggregate known as the kinetoplast that duplicates asynchronously from the nucleus[1,19]. A single flagellum nucleates from a basal body docked to an invagination of the cell surface known as the flagellar pocket, which in some life-cycle stages serves as a key endocytic and exocytic compartment that is crucial to host immune evasion [9,20]. The flagellum traverses the pocket and then emerges onto the cell surface. In some cases, the flagellum is attached laterally along the cell surface via a series of junctional complexes known as the flagellum attachment zone (FAZ), while in others the flagellum is detached from the cell surface [21,22]. Cells with free flagella still retain a small FAZ segment near the neck of the flagellar pocket (**Fig 1**) [23]. A subset of trypanosomatids have a separate tube-shaped endocytic membrane compartment flanked by a set of cytoplasmic microtubules with an opening next to the flagellar pocket, known as the cytostome-cytopharynx[24,25]. Nearly all trypanosomatids have a set of subpellicular microtubules underlying the plasma membrane that persists throughout the cell cycle and is responsible for shaping the cell body [26,27]. These microtubules are heavily crosslinked to each other and to the plasma membrane.

**Fig 1.**
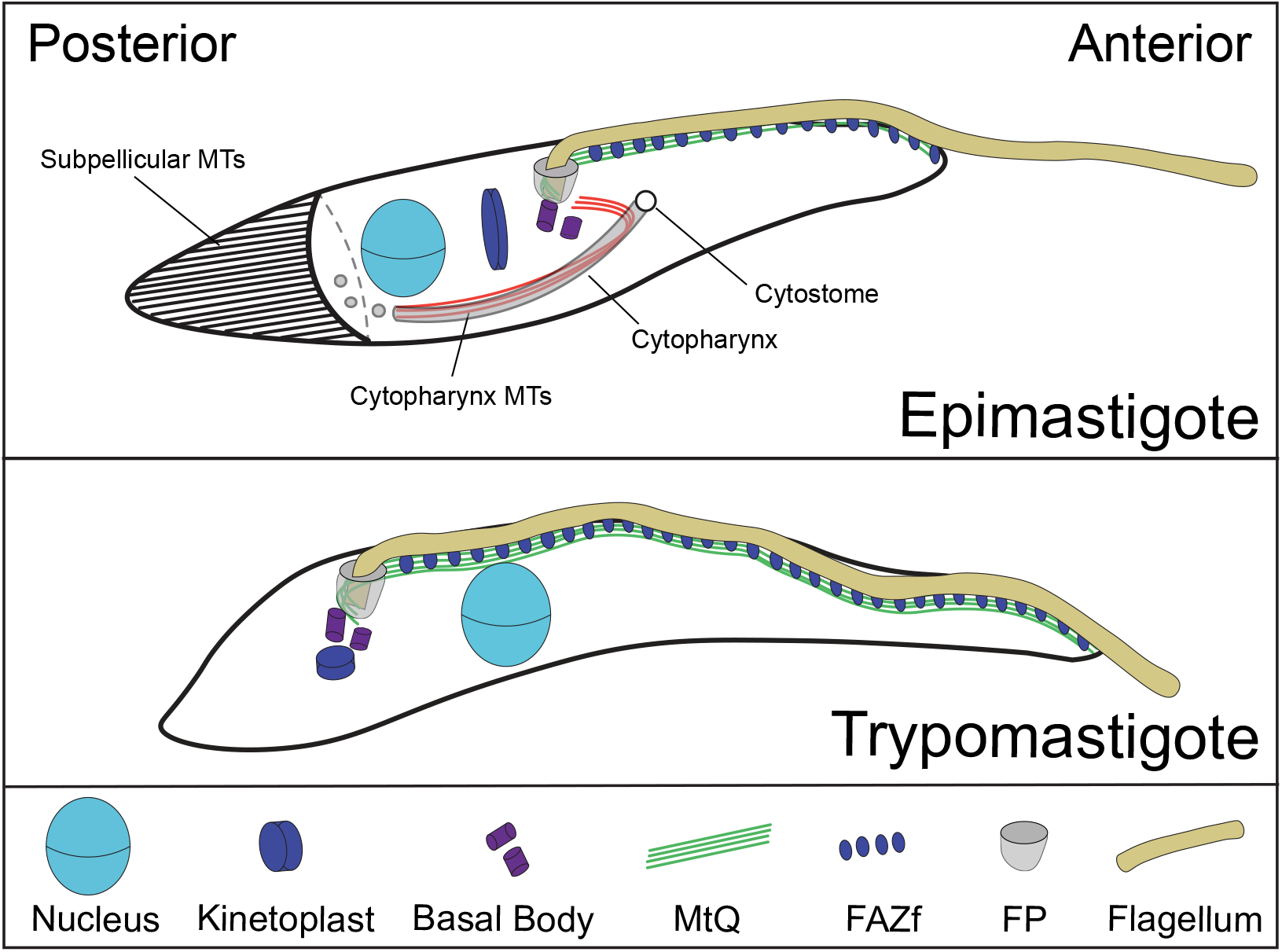
The morphotypes of Trypanosoma cruzi. All *T. cruzi* morphotypes contain a flagellum that is nucleated from a basal body which is docked to the membrane of the flagellar pocket. The basal body is also connected to the kinetoplast, which is the mitochondrial DNA aggregate. The shape of the cell bodies is defined by a set of subpellicular MTs that underlie the plasma membrane (depicted only in the epimastigote form). In the epimastigote and trypomastigote forms, the flagellum emerges onto the cell surface and is attached to the cell body by the flagellum attachment zone (comprising the MtQ and FAZ filament), both of which extend towards the anterior end of the cell body. The primary morphologic difference between epimastigotes and trypomastigotes is the positioning of the kinetoplast-basal body-flagellar pocket, which is posterior to the nucleus in epimastigotes and anterior in trypomastigotes.

Trypanosomatids assume a wide range of cell morphologies as they cycle within their host environments. In the bloodstream of their vertebrate hosts, many trypanosomatids assume a trypomastigote morphology, with the kinetoplast, flagellum, and flagellar pocket on the posterior side of the nucleus, near the cell posterior [1,19]. The flagellum is attached to the plasma membrane by an extended FAZ. It has been proposed that this morphology is optimized for motility in crowded, high-viscosity solutions such as blood [28,29]. *T. brucei* and *T. cruzi* bloodstream forms assume a trypomastigote morphology, while *Leishmania* assume a promastigote morphology, where the kinetoplast, flagellar pocket, and minimally attached flagellum are on the anterior side of the nucleus [30]. At certain locations within the insect host, *T. brucei* and *T. cruzi* adopt an epimastigote morphology, with the kinetoplast and flagellum positioned on the anterior side of the nucleus and a segment of attached flagellum extending towards the anterior end of the cell [31–33]. This morphology is thought to be optimal for attachment to epithelial surfaces, which is a common part of the insect infection stages (**Fig 1**). Trypanosomatids remain polarized throughout their cell cycle, which has required morphotypes to adapt the mechanisms used for duplicating and positioning organelles during cell division to suit their specific needs [7].

In this work, we have studied the cell division of *T. cruzi* epimastigotes using quantitative morphometrics and epitope-tagged proteins that are important cell cycle components in *T. brucei*. We show that the epimastigote cell body widens and shortens as it divides, suggesting that the cell division mechanism in this form is more similar to the process in *Lieshmania* promastigotes. Epimastigote cell division appears highly asymmetric, as the daughter cell inheriting the new flagellum is smaller than the old-flagellum daughter. Using live-cell imaging, we show that the rate of daughter cell divisions differs by nearly 5 h, which suggests that the asymmetry may result in one daughter cell taking longer to complete a subsequent round of division. The construction of a new posterior end recruits many of the cell cycle regulated proteins seen at the tip of the new FAZ in *T. brucei*, along with tubulin posttranslational modifications usually confined to the FAZ. These results provide a framework for describing the cell division mechanism in the *T. cruzi* epimastigote, which is a common morphology found in many trypanosomatids that has not been studied in detail previously.

## Results

The cell division mechanism trypanosomatids employ is influenced by the arrangement of the organelles within their specific morphotype [7]. The maturation of the probasal body and assembly of the new flagellum are usually the earliest events, followed by the assembly of a new FAZ, the duplication of the pocket and kinetoplast, and finally nuclear duplication [10,34]. The subpellicular array must also be enlarged to accommodate the duplicating organelles and to situate them within the cell body so that cytokinesis will produce two daughter cells with the correct complement and arrangement of organelles [35]. Cleavage furrow ingression initiates from the anterior end of the cell body and moves towards the posterior end to complete cell division. In *T. brucei* trypomastigotes, lengthening of the cell body is the primary means of accommodating duplicated organelles and positioning them for inheritance; *Leishmania* promastigotes widen their cell bodies while simultaneously shortening them [36,37]. It has been speculated that the extended segment of attached flagellum in trypomastigotes may limit their ability to widen their cell body, which has led to the lengthening mechanism [7]. The *T. cruzi* epimastigote form, which has an attached flagellum like trypomastigotes but an anterior-located flagellar pocket and kinetoplast, could provide insight into the effect of flagellar attachment on cell division.

We employed Y-strain epimastigotes constitutively expressing *Streptococcus pyogenes* Cas9 and T7 polymerase to tag genes at their native loci with three copies of the Ty1 epitope fused to mNeonGreen, which allowed us to observe the native fluorescence of the tagged protein or to employ anti-Ty1 antibodies for detection [14,17,38–40]. We selected marker proteins based on recent work in *T. brucei* and *Leishmania* that identified “landmark” proteins that provide bright signal as mNeonGreen fusions for a range of cellular compartments [41]. For each protein, we tagged the terminus that yielded the brightest signal based on the TrypTag whole-genome tagging database [42]. The Cas9-T7 expressing epimastigotes were nucleofected with guide DNAs containing a T7 promoter to produce a guide RNA targeting one terminus of the gene of interest and a healing template that introduces the 3X-Ty1-mNeonGreen tag, an intergenic region, and a selection marker. We tagged a series of proteins that localize to the FAZ, extending new FAZ, basal body, and cell posterior in *T. brucei* trypomastigotes to determine if their localization was similar in *T. cruzi* epimastigotes and to use these marker proteins to study the mechanism of cell division in this morphotype.

We first observed the FAZ using 3X-Ty1-mNeonGreen::FAZ25 (TcYC6_0125610) and the monoclonal antibody 1B41, which labels the FAZ in *T. brucei*. The 1B41 antibody is thought to label a novel post-translational modification (PTM) of beta tubulin present in the segment of the MtQ that is adjacent to the FAZ filament [43,44]. We noted that FAZ25 was present in a single thin structure underlying the flagellum, as would be expected for the FAZ in epimastigote-form *T. cruzi* (**Fig 2**) [45]. The 1B41 signal generally colocalized with FAZ25, but we observed additional 1B41-positive structures in dividing cells. Early cell cycle cells containing one **N**ucleus and one **K**inetoplast (1N1K) could be separated into two populations: smaller cells with a short FAZ and a separate 1B41 focus at the cell posterior (**Fig 2A**), and larger cells with a longer FAZ and no posterior 1B41 labeling or a much less intense signal than the shorter 1N1K cells (**Fig 2B**). Upon initiation of new FAZ assembly, a small structure was observed in both the FAZ25 and 1B41 channels on the ventral side of the old FAZ. The 1B41 punctum at the cell posterior, which was frequently much brighter than the 1B41 signal at both the old and new FAZ, became visible at this point (**Fig 2C, 2D**). In 2N2K cells with ingressing cleavage furrows, the posterior 1B41 punctum marked the last point of connection between the daughter cells (**Fig 2E**). The new-flagellum daughter appeared to be significantly smaller than the old-flagellum daughter in cells that were about to complete cytokinesis. Because of this asymmetry, we propose that the small 1N1K cells with the bright 1B41 puncta are likely to be new-flagellum daughter cells that have just completed cytokinesis. These smaller cells do not appear to enter cell division in their current state, which suggests that they grow in size and lose their 1B41 puncta over time prior to initiating cell division. The longer 1N1K cells lacking the 1B41 puncta are likely to be a mixture of the old-flagellum daughter cells and the new-flagellum daughter cells that have undergone subsequent growth after the completion of cell division. This would also suggest that the 1B41 punctum is inherited asymmetrically, with the new-flagellum daughter cell retaining the bulk of the signal (**Fig 3A**). This asymmetric inheritance is similar to what is observed in the partitioning of the flagella connector in procyclic *T. brucei* cells, which allowed the definitive identification of the new- and old-flagellum daughter cells after the completion of cell division [46].

**Fig 2.**
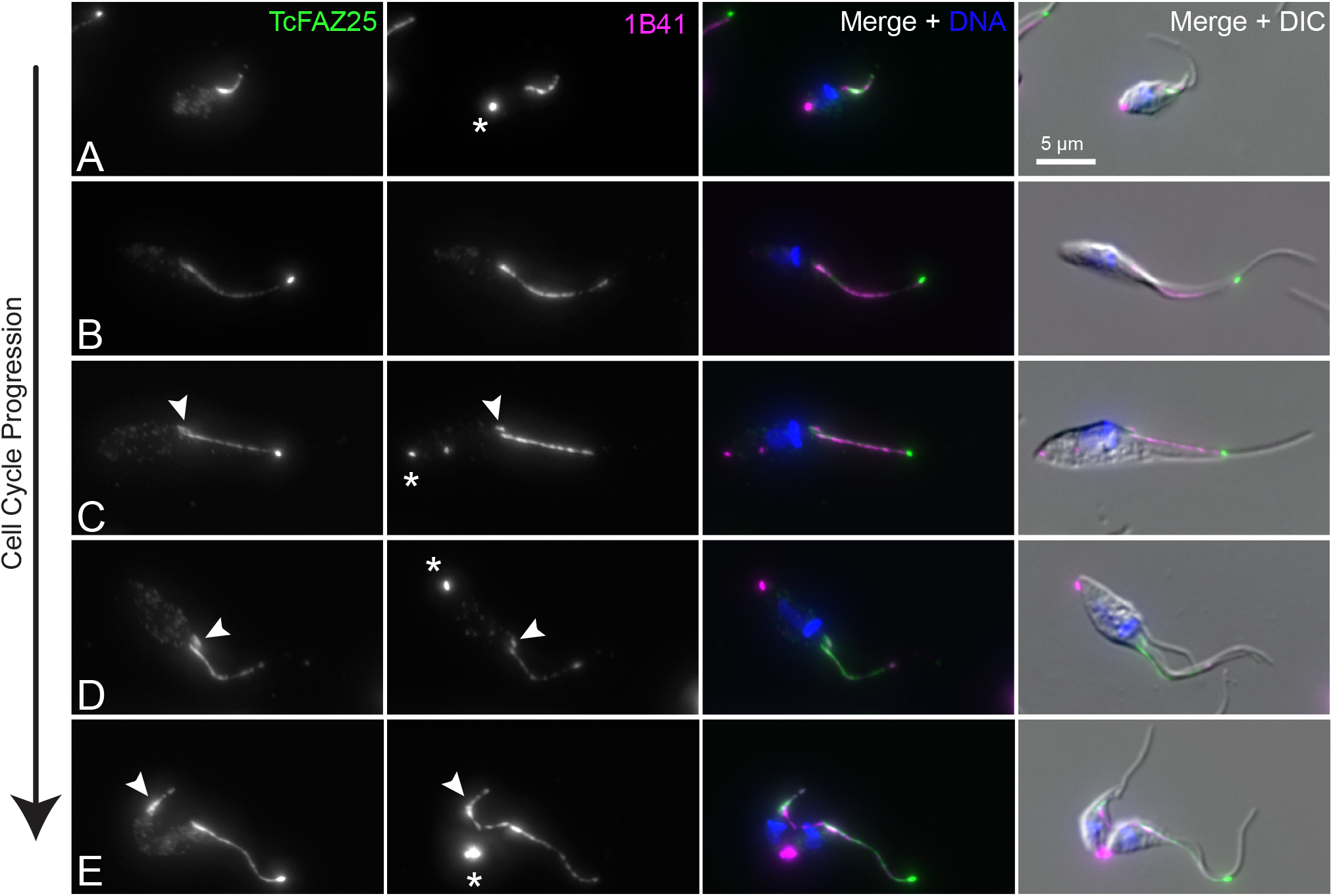
*T. cruzi* epimastigotes incorporate tubulin PTMs at the cell posterior during cell division. Epimastigote cells carrying a 3xTy1-mNeonGreen::TcFAZ25 allele were fixed and labeled with 1B41 to label the FAZ and posterior punctum (1B41, magenta), anti-Ty1 to label FAZ25 (TcFAZ25, green), and DAPI to label DNA (DNA, blue). Cells were then imaged using epifluorescence and DIC microscopy. (**A**) Very small 1N1K cells carried a 1B41 punctum at their cell posterior, which was much weaker or absent in larger 1N1K cells (**B**). (**C**) The new FAZ was appeared on the dorsal side of the cell and labeled was FAZ25 and 1B41 positive. 1B41 signal appeared at the posterior end at this point. (**D**) As the new FAZ and flagellum extended, the posterior 1B41 punctum increased in brightness. (**E**) Once the cell has begun cytokinesis, the 1B41 punctum is present at the extreme posterior of the cell body. The new-flagellum daughter cell appears to inherit a much shorter new FAZ and flagellum than the old-flagellum daughter. Asterisks highlight the 1B41 posterior punctum labeling, while the arrowheads identify the new FAZ.

**Fig 3.**
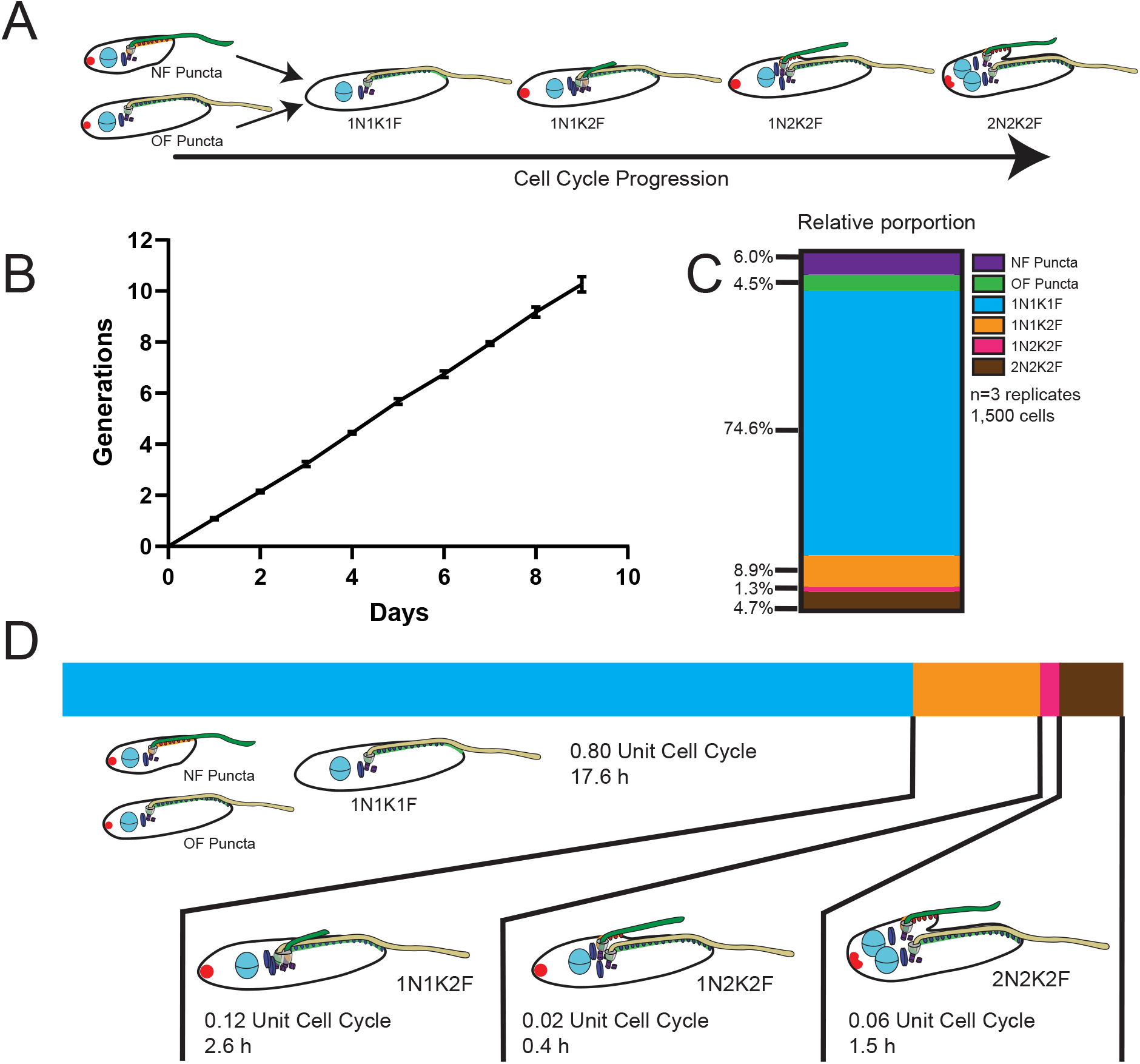
Defining and quantifying morphogenic intermediates in *T. cruzi* epimastigote cell division. (**A**) Morphogenic intermediates were defined by the presence of a 1B41 punctum and if they were the new flagellum or old flagellum daughter (new-flagellum puncta: NF Puncta; old-flagellum puncta: OF puncta), or their nuclear, kinetoplast, and flagellum state (xNxKxF). Note that all cells that have begun to flagellar duplication are labeled with a 1B41 punctum. (**B**) Cell counts over 9 d were used to calculate the cell division time of the Cas9-T7 epimastigote cells line. (**C**) Graph depicting the percentages of cells in each state as defined in (**A**). (**D**) Using the length of the cell cycle as measured in (**B**) and the percentages of cells in different cell cycle states measured in (**C**) allows the calculation of the time that the parasites spend in each state that was defined in (**A**).

The novel localization pattern of the 1B41 antibody in *T. cruzi* suggests that the cell posterior is transiently modified with a tubulin PTM during cell division that is usually confined to the MtQ segment adjacent to the FAZ filament *in T. brucei*. The posterior end contains many of the plus ends of the subpellicular microtubules, which could be specifically targeted by PTMs as part of the construction of the new cell posterior during cell division [26]. We labeled *T. cruzi* with antibodies that detect tubulin tyrosination, acetylation, and two forms of glutamylation to determine if they have similar labeling patterns to 1B41 in dividing cells, which could indicate that they are labeling the same tubulin PTM (**S1 Fig**). The YL1/2 antibody, which detects tyrosinated tubulin that is added to growing microtubules, labeled the cell posterior, the posterior part of the subpellicular array, and the basal body [47–49]. The GT335 antibody, which labels polyglutamate chains of all lengths, and IN105, which labels polyglutamate chains at least three units in length, both labeled the entirety of the subpellicular array, including the flagellum [50,51]. The 6-11B-1 antibody, which labels acetylated tubulin, labeled the entirety of the array and the flagellum [52]. Considering the broad distribution of these antibody signals across multiple structures that lack 1B41 signal, it is likely that the 1B41 epitope is not attributable to tubulin tyrosination, acetylation, or glutamylation.

We performed a quantitative analysis of the cell cycle stages we established using the localization patterns of FAZ25 and 1B41 to measure how *T. cruzi* changes as it divides. We employed a statistical method that relies on an asynchronously dividing culture to establish the length of time that cells spend in different states [53,54]. This approach relies on a precise measurement of parasite doubling time, which we assessed for our Cas9-T7 cell line over the course of 9 d. We found a persistent doubling time of 22.1 h, which we used along with the number of cells in each of our FAZ25/1B41 categories to define the epimastigote cell cycle (**Fig 3B and 3C**). *T. cruzi* epimastigotes spend 80% of their time as one **<u>N</u>**ucleus, one **<u>K</u>**inetoplast, and one **<u>F</u>**lagellum (1N1K1F) cells, representing 17.6 h of the cell cycle. New- and old-flagellum daughter cells containing the 1B41 posterior puncta account for 6% and 4.4% of the cell population, respectively (**Fig 3C**). New flagellum and FAZ growth initiates from a point on the ventral side of the cell next to the existing FAZ and flagellum. The 1B41 puncta appears at this time as well. The briefest window of the cell cycle occurs after kinetoplast duplication and prior to nuclear duplication (1N2K), which represents just under 30 min. It should be noted that at this stage the *T. cruzi* kinetoplasts label brightly with DAPI and are in close proximity to one another, making it difficult to clearly determine when the cells reach the 2K state. Consequently, the number of cells we identified as 1N2K2F, and therefore the length of time the cells spend in this state, is likely an underestimate. *T. cruzi* spends considerably longer in the 2N2K2F stage; almost all of these cells have an ingressing cleavage furrow, which suggests that cytokinesis initiates rapidly after the completion of karyokinesis (**Fig 3D**).

We also monitored the morphologic changes that occur in epimastigotes as they divide. We measured the width of the cell along its shortest axis and the overall cell length including the extension of the flagellum and the cell body alone. We also measured the distances between the cell posterior and the kinetoplast and nucleus, along with the lengths of the flagellum and FAZ (**Fig 4A**). We noted that the cell body length and the distances between the nucleus and kinetoplast and the posterior end of the cell decline towards the end of cell division while the overall width of the cell body increases (**Fig 4B-E, 4H**). The cell body shortening seems to be restricted to the cell posterior as the old FAZ and flagellum lengths remain unchanged (**Fig 4F and 4E**). The cells that we previously identified as small with a bright 1B41 punctum at their cell posteriors were shorter than 1N1K1F cells lacking puncta or with weak puncta labeling, which suggests that the new-flagellum daughter cells are significantly shorter than the old-flagellum daughter cells right after completing cytokinesis.

**Fig 4.**
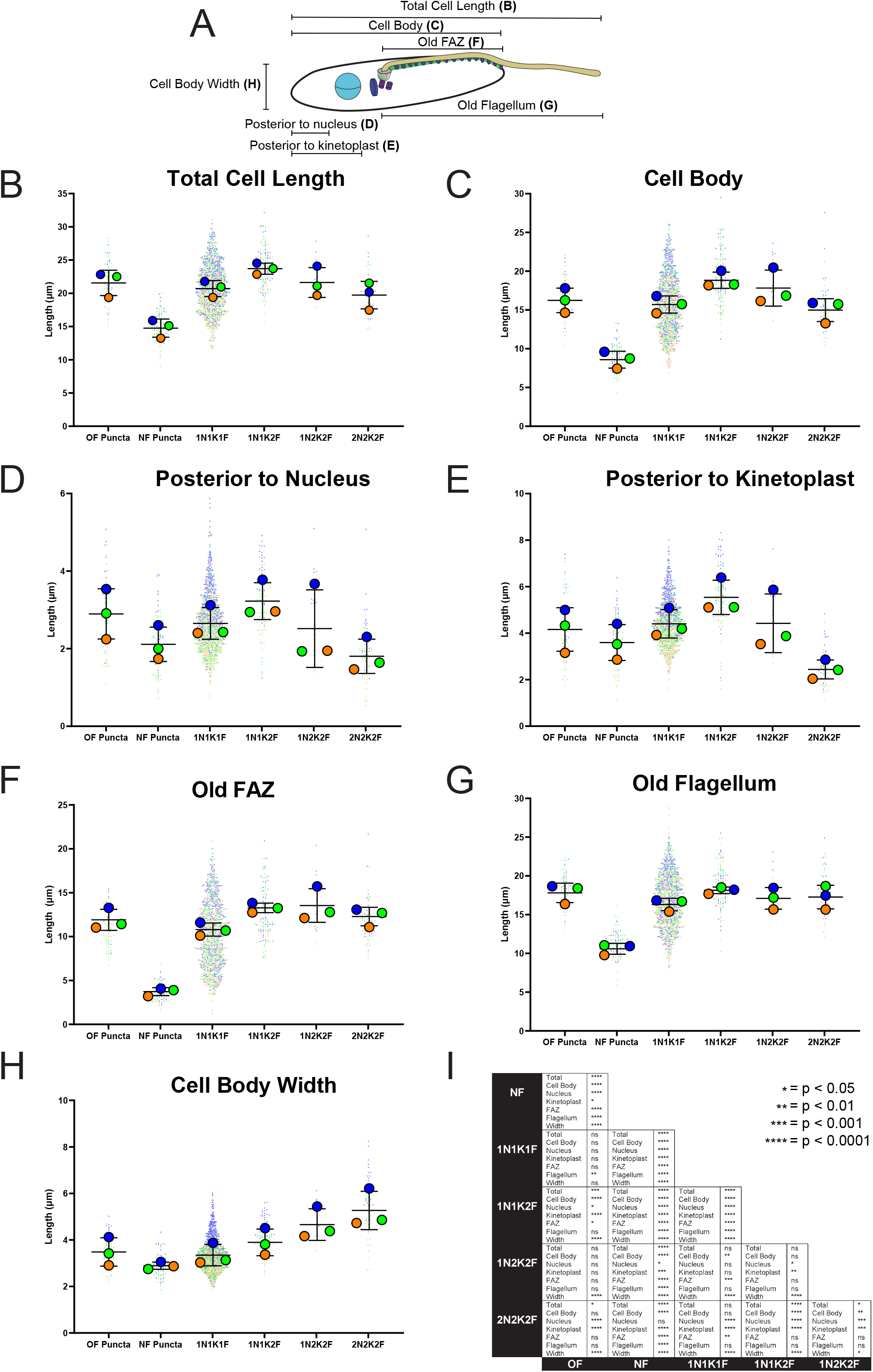
Quantitation of changes to cell morphology over the course of *T. cruzi* epimastigote cell division. (**A**) Cells were fixed and stained with 1B41 to label the FAZ and posterior punctum and DAPI to label DNA. The cells were then imaged using epifluorescence and DIC microscopy, followed by measurement of all the cell features depicted in this schematic. All graphs were generated as SuperPlots, where each independent experiment is shown in a separate color, with the large circles represent the mean of each independent experiment, and black lines represent the mean and the standard deviation. (**B**) Measurements of total cell lengths, which includes flagellar overhang, broken down into the cell division categories defined in **Fig 3A**. (**C**) Measurements of cell body lengths, broken down into the cell division categories defined in **Fig 3A**. (**D**) Measurements of the distance between the cell posterior and the nucleus, broken down into the cell division categories defined in **Fig 3A**. (**E**) Measurements of the distance between the cell posterior to the kinetoplast, broken down into the cell division categories defined in **Fig 3A**. (**F**) Measurements of the length of the old FAZ, broken down into the cell division categories defined in **Fig 3A**. (**G**) Measurements of the length of the old flagellum, broken down into the cell division categories defined in **Fig 3A**. (**H**) Measurements of the width of the cell body, broken down into the cell division categories defined in **Fig 3A**. (**I**) Table showing significance levels between each pair-wise comparison. Statistical analysis was done by one-way ANOVA. ns = not significant, * p < 0.05, ** p < 0.01, ***p < 0.001, ****p < 0.0001.

We focused on the difference between the new and old FAZ and flagellum lengths as an indicator of the asymmetry present during the late stages of cell division (**Fig 5A**). The new FAZ averages 2.2 microns in cells that are undergoing cytokinesis, while the old FAZ remains approximately 12 microns long throughout cell division, showing the extreme asymmetry between the inherited structures (**Fig 4F and 5B**). The difference between the length of the old and new flagella was smaller, with new flagellum lengths at the latest stages of cell division reaching 9 microns, compared to 18 microns for the old flagellum (**Fig 4G and 5C**). This result strongly suggests that most of the asymmetry observed during *T. cruzi* epimastigote cell division is attributable to the construction of a smaller new cell body that is inherited by the new flagellum/FAZ daughter. The new flagellum daughter cell must then increase its cell body length and extend the FAZ prior to initiating another round of cell division (**Fig 5D and 5E**).

**Fig 5.**
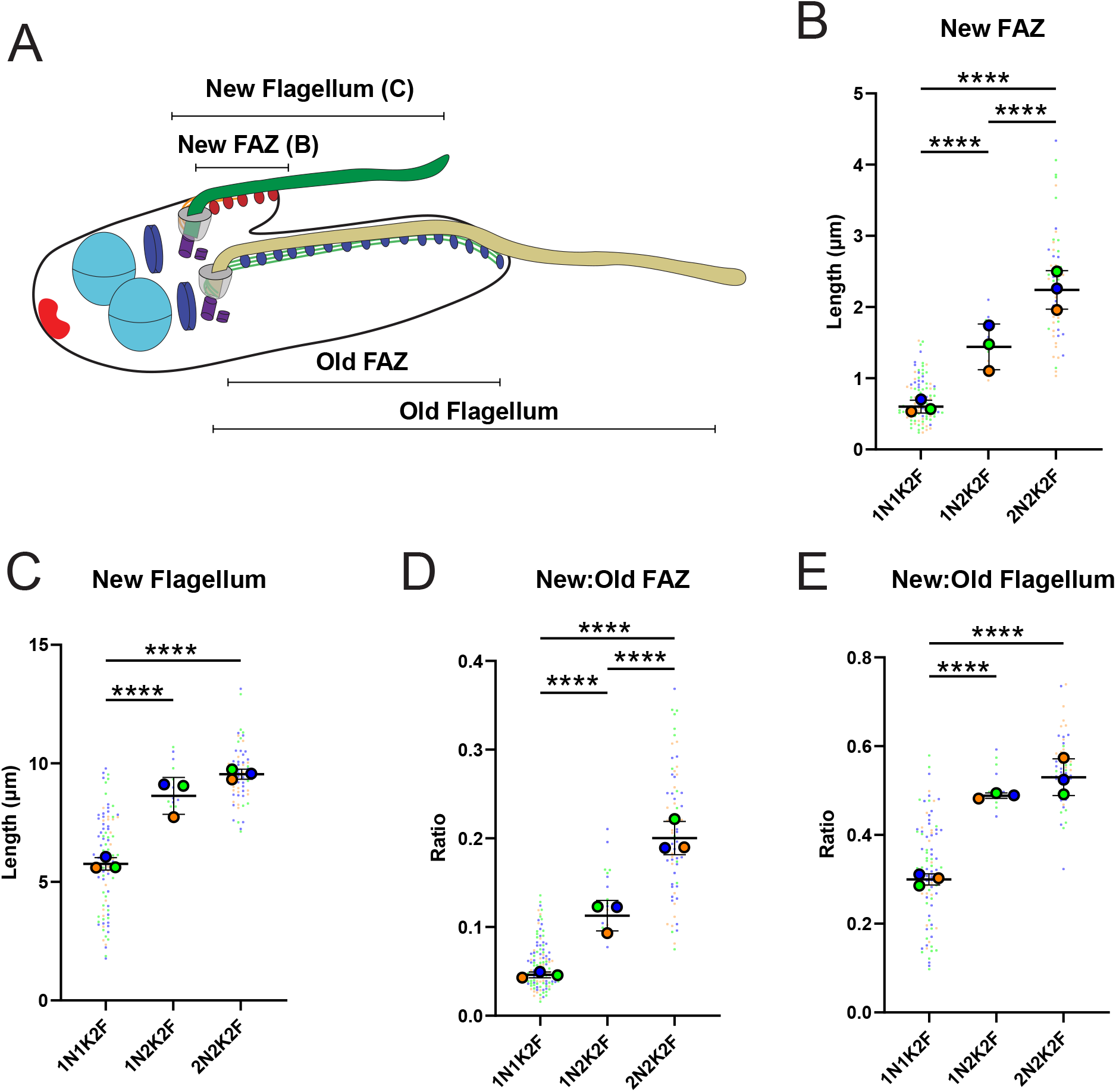
The new FAZ does not reach length parity with the old FAZ prior to the completion of cell division. (**A**) Cells were fixed and stained with 1B41 to label the FAZ and posterior punctum and DAPI to label DNA. The cells were then imaged using epifluorescence and DIC microscopy, followed by measurement of all the cell features depicted in this schematic. All graphs were generated as SuperPlots, where each independent experiment is shown in a separate color, with the large circles represent the mean of each independent experiment, and black lines represent the mean and the standard deviation. (**B**) Measurements of the length of the new FAZ as cells progress through cell division. (**C**) Measurements of the length of the new flagellum as cells progress through cell division. (**D**) Ratio of the length of the new FAZ to the old FAZ as cells progress through cell division. (**E**) Ratio of the length of the new flagellum to the old flagellum as cells progress through cell division. Statistical analysis was done by one-way ANOVA ****p < 0.0001.

We labeled the parental Y-strain *T. cruzi* line with 1B41 and quantitated its cell morphology to confirm that the posterior punctum 1B41 signal and the asymmetric cell division we observed was not due to the presence of Cas9 and T7 polymerase in our tagging cell line. We found that the parental strain also has the posterior 1B41 signal and widens and shortens over the course of cell division (**S2 Fig**). We also noted the asymmetry between the new flagellum and old flagellum daughter cells during the late stages of cytokinesis, and that 1N1K cells with the posterior 1B41 punctum were significantly smaller than 1N1K lacking the punctum. We also labeled Brazil A4 strain *T. cruzi* epimastigotes with 1B41 and saw the posterior punctum and asymmetry in the daughter cells during division, which suggests that these features are generally conserved among different *T. cruzi* strains.

We next tagged the protein SAS-6 (TcYC6_0070700) to gain insight into the basal body duplication cycle in *T. cruzi*. In *T. brucei*, TbSAS-6 is present at both the pro and mature basal body throughout cell division [55,56]. We tagged TcSAS-6 with 3X-Ty1-mNeonGreen and followed its localization over the cell cycle with 1B41 as a colabel (**Fig 6**). TcSAS-6 labels both the pro and mature basal body in 1N1K1F cells containing the posterior 1B41 puncta, denoting cells that had just recently completed cell division. Curiously, we noted in 1N1K cells that had produced a new flagellum (arrowhead) and were developing a new 1B41 punctum still only had two TcSAS-6-positive structures. In *T. brucei*, cells with two flagella have four TbSAS-6-positive structures, reflecting two pairs of pro and mature basal bodies. At later points in cell division, we noted cells with 3 and 4 TcSAS-6-positive structures, suggesting that SAS-6 is eventually incorporated into all of the basal bodies (**Fig 6A**). We used the pan-centrin antibody 20H5 as another marker that should persistently label both the pro and mature basal body [57]. In cells colabeled for TcSAS-6, we were able to observe 20H5-positive basal bodies at the base of the flagella in dividing cells that lacked SAS-6 labeling, showing that the absence of TcSAS-6 did not impact other persistent basal body markers (**S3A Fig**). As another control, we looked at the intrinsic fluorescence of the mNeonGreen appended to TcSAS-6 to make sure that antibody accessibility was not limiting our ability to detect all the SAS-6 present in the cell. The mNeonGreen fluorescence followed the same pattern as observed by immunofluorescence (**S3B Fig**).

**Fig 6.**
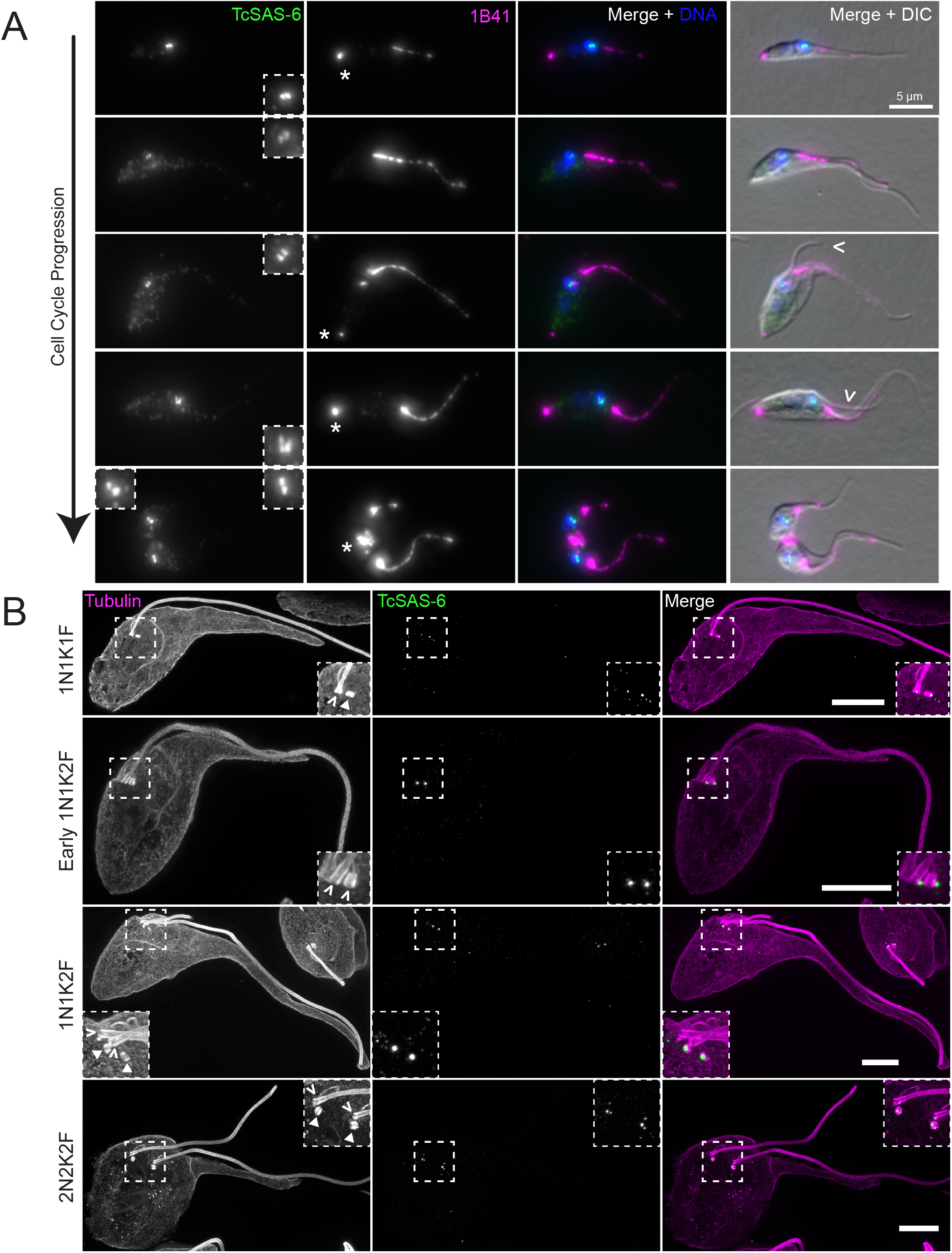
TcSAS-6 levels fluctuate on the basal body during cell division. (**A**) Cells carrying a 3xTy1-mNeonGreen::TcSAS-6 allele were fixed and labeled with 1B41 (1B41, magenta) to label the FAZ and posterior punctum, anti-Ty1 (TcSAS-6, green) to label the basal body, and DAPI to label DNA (DNA, blue). The cells were then imaged using epifluorescence and DIC microscopy. Inserts show a 3X enlargement of the basal body region. The empty arrowheads identify the new flagellum; the asterisks highlight the 1B41 posterior punctum. (**B**) Cells carrying a 3xTy1-mNeonGreen::TcSAS-6 allele were fixed and processed for U-ExM. Samples were labeled with anti-Ty1 (TcSAS-6, green) to label the basal body and TAT1 anti-tubulin antibody (Tubulin, magenta) to label the tubulin-containing structures within the cell. The samples were then imaged using SoRa superresolution microscopy. Boxed insets are a 3X magnification of the basal body region. The empty arrowheads identify the mature basal body, while the filled arrowheads identify the probasal body. Scale bars: 10 µm.

We employed ultrastructure expansion microscopy (U-ExM) coupled with SoRa super-resolution imaging to better characterize the localization pattern of TcSAS-6 in *T. cruzi* (**Fig 6B**) [58–60]. U-ExM provides an approximately 5-fold enhancement in resolution that is further enhanced by √2 past the light limit with SoRA, which also provides confocal sectioning [61]. We used anti-Ty1 labeling for TcSAS-6 and the TAT1 anti-tubulin antibody to label the flagellum, basal body, the microtubules associated with the cytostome, and subpellicular array [62]. In the expanded samples, we were able to clearly identify pro and mature basal bodies at all stages of cell division. We observed TcSAS-6 labeling of both the pro and mature basal body in cells with a single flagellum. In cells that had just begun to duplicate the basal body, which can be identified by the rotation of the maturing probasal body to facilitate docking with the flagellar pocket membrane, the two TcSAS-6 puncta were no longer associated with the barrel of the pro or mature basal body, but appeared to be adjacent in two discrete foci. There were still two SAS-6 puncta associated with the duplicated basal bodies in cells that had begun to divide that contained a short new flagellum, but the puncta were found exclusively on both probasal bodies. There was no TcSAS-6 signal on the mature basal bodies that nucleated the old and new flagella. At later stages of cell division, which we could identify by the increased length of the new flagellum, we could observe SAS-6 signal restored to the mature basal bodies, which is consistent with our immunofluorescence results (**Fig 6B**). With the additional resolution provided by U-ExM and SoRa imaging, we noted that the TcSAS-6 signal on the probasal body is actually two distinct puncta at this stage of cell division, one that appears to be associated with the base of the probasal body and another punctum on the side of the probasal body at the approximate midpoint of the structure. These results suggest that SAS-6 levels decline significantly at the mature basal body in *T. cruzi* during cell division and that this decline may be linked to basal body duplication.

Proteins that transiently localize to the tip of the extending new FAZ play an essential role in positioning the cleavage furrow in *T. brucei* trypomastigotes [10]. Several cytokinetic proteins are present at the tip of the new FAZ in *T. brucei* trypomastigotes and are thought to use its extension and position within the cell body to guide the placement of the new cell anterior and designate the starting point for cleavage furrow ingression. Among these proteins is the cytokinetic scaffolding protein TbTOEFAZ1 (also known as TbCIF1), which is essential for cleavage furrow placement in *T. brucei* cells [63,64]. We tagged the *T. cruzi* homolog of TcTOEFAZ1 (TcYC6_0051010) and monitored its localization. In *T. brucei*, TbTOEFAZ1 expression is found at the tip of the extending new FAZ and then along the cleavage furrow. In *T. cruzi*, we noted that all cells, regardless of cell cycle stage, expressed TcTOEFAZ1 (**Fig 7**). In the early stages of the cell cycle prior to new FAZ assembly, TcTOEFAZ1 was localized to a short bar-like structure adjacent to the proximal end of the old FAZ. This localization is similar to the position of the new FAZ that appears during the later stages of cell division. In cells that had begun to assemble the new FAZ, TcTOEFAZ1 localized along the length of the new structure, but did not form a focus at the tip as it does in *T. brucei*. TcTOEFAZ1 is also present at the point where the cleavage furrow ingresses in *T. cruzi*, which is found between the two flagella (**Fig 7A**).

**Fig 7.**
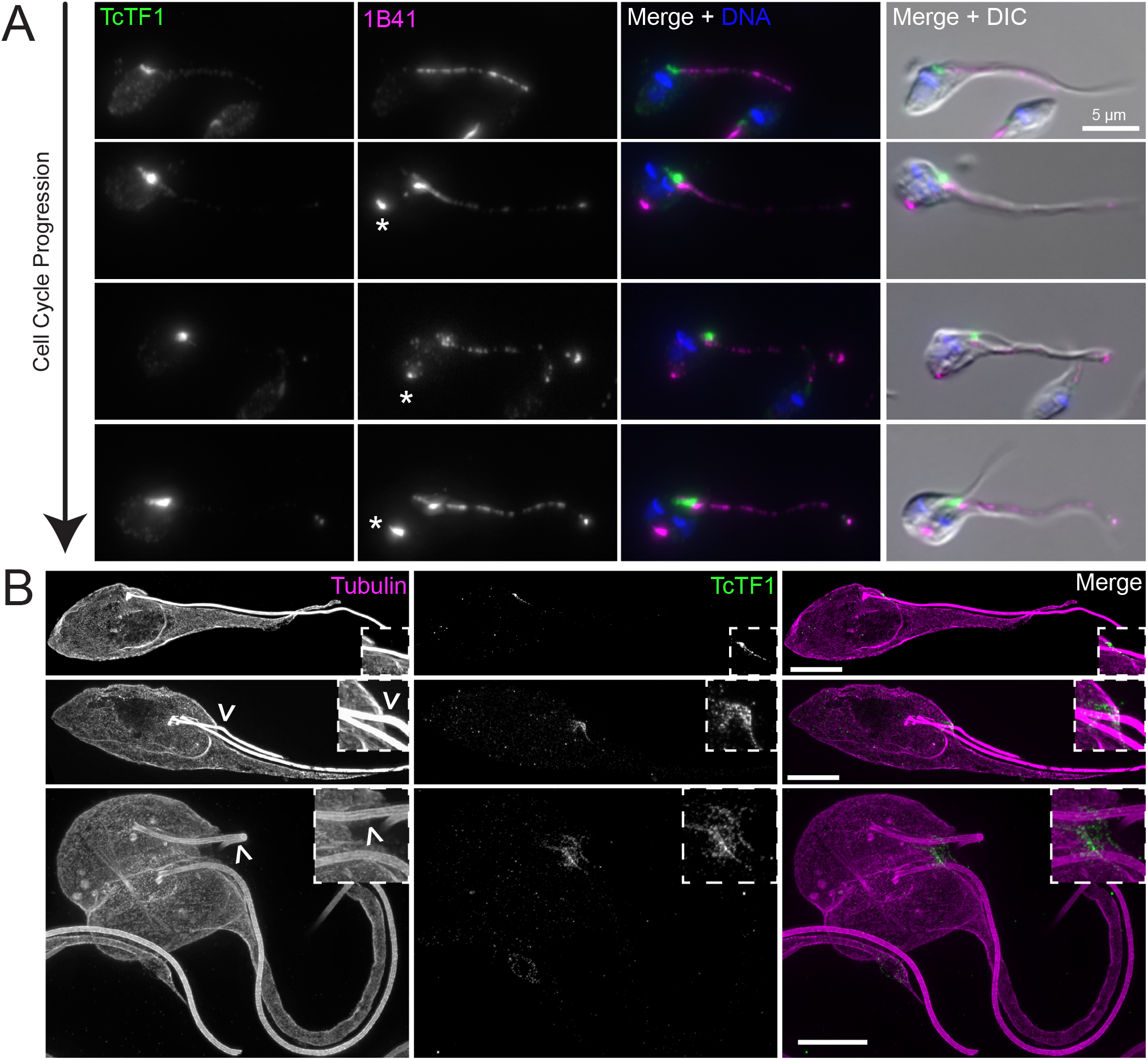
TcTOEFAZ1 localizes to the ventral side of the flagellum is expressed during all cell stages. (**A**) Cells carrying a 3xTy1-mNeonGreen::TcTOEFAZ1 allele were fixed and labeled with 1B41 to label the FAZ and posterior punctum (1B41, magenta), anti-Ty1 to label TcTOEFAZ1 (TcTF1, green), and DAPI to label DNA (DNA, blue). The cells were then imaged using fluorescence and DIC microscopy. The asterisks identify the 1B41 posterior punctum. (**B**) Cells carrying a 3xTy1-mNeonGreen::TcTOEFAZ1 allele were fixed and processed for U-ExM. Samples were labeled with anti-Ty1 to label TOEFAZ1 (TcTF1, green) and TAT1 anti-tubulin antibody to label the tubulin-containing structures within the cell (Tubulin, magenta). The samples were then imaged using SoRa super-resolution microscopy. Boxed insets are a 3X magnification of the basal body region. The empty arrowheads point toward the new flagellum. Scale bars: 10 µm.

We employed U-ExM and SoRa imaging to further establish the localization of TcTOEFAZ1, using anti-tubulin as a colabel (**Fig 7B**). In early cell cycle cells, TcTOEFAZ1 localized to a structure on the ventral side of the new flagellum, along with a small amount of labeling underlying the segment of the old flagellum just as it exits the flagellar pocket. In dividing cells, TcTOEFAZ1 localization appeared to encircle the point where the new flagellum exits the flagellar pocket onto the cell surface. TcTOEFAZ1 was present at the position of the ingressing furrow, in the area between the new and old flagellum. We were unable to directly compare TcTOEFAZ1 localization to a FAZ marker because U-ExM imaging does not appear to preserve sufficient signal for 1B41 or FAZ25.

The Polo-like kinase (TbPLK) homolog present in *T. brucei* binds to and phosphorylates TbTOEFAZ1, which is essential for TbPLK localization to the tip of the extending FAZ [65,66]. Recently, a second homolog of TbPLK has been identified in *T. brucei*, but little is known about its function [67]. We tagged the TbPLK1 homolog in *T. cruzi* to compare its localization pattern to the *T. brucei* homolog (**Fig 8A**). TcPLK1 (TcYC6_0047790) was not expressed in cells early in the cell cycle, which is consistent with TbPLK expression patterns in *T. brucei*. During the early stages of cell division TcPLK1 initially localized to the basal body, as is the case during *T. brucei* cell division [68]. However, the kinase does not subsequently collect at the tip of the new FAZ, but rather localizes along the length of the old FAZ. TcPLK1 also localizes to the new FAZ once this structure becomes visible. TcPLK1 is subsequently present at the cell posterior, appearing just prior to the 1B41-positive puncta, then colocalizing with it, which represents a novel localization pattern for the kinase in trypanosomatids. TcPLK1 does not appear to track along the cleavage furrow during the latest stages of cell division, which is consistent with its localization in *T. brucei* trypomastigotes (**Fig 8A**).

**Fig 8.**
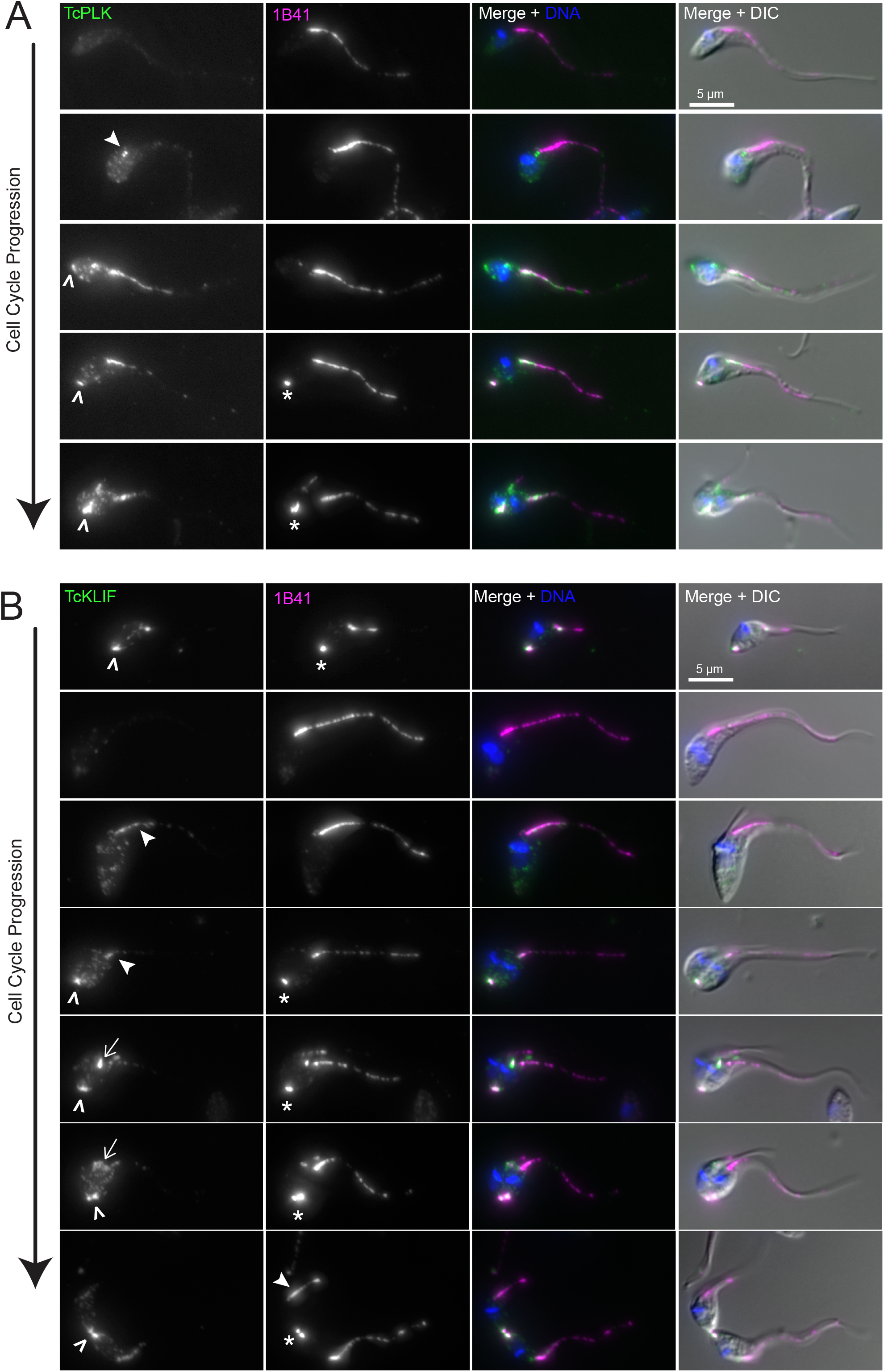
TcPLK and TcKLIF localize to the FAZ and the posterior punctum during cell division. (**A**) Cells carrying a 3xTy1-mNeonGreen::TcPLK allele were fixed and labeled with 1B41 to label the FAZ and posterior punctum (1B41, magenta), anti-Ty1 to label TcPLK (TcPLK, green), and DAPI to label DNA (DNA, blue). The cells were then imaged using fluorescence and DIC microscopy. Asterisks identify the 1B41 posterior punctum, empty arrowheads show TcPLK recruitment to the cell posterior, and the filled arrowhead shows TcPLK localization to the basal body. (**B**) Cells carrying a 3xTy1-mNeonGreen::TcKLIF allele were fixed and labeled with 1B41 to label the FAZ and posterior punctum (1B41, magenta), anti-Ty1 to label TcKLIF (TcKLIF, green), and DAPI to label DNA (DNA, blue). Asterisks show the 1B41 posterior punctum, empty arrowheads show TcKLIF location to the cell posterior, filled arrowheads show TcKLIF localization to the FAZ, and arrows show TcKLIF localization to the ingressing furrow.

In *T. brucei,* the orphan kinesin TbKLIF plays an important role in the last stages of cell division during the construction of a new cell posterior [69–71]. In *T. cruzi*, 3X-Ty1-mNeonGreen::TcKLIF (TcYC6_0055740) colocalized with the 1B41 posterior puncta in cells that had recently completed cell division (**Fig 8B**). 1N1K cells lacking the puncta no longer had KLIF signal, suggesting that KLIF is downregulated during the early stages of the cell cycle. Once the cells had begun to divide, we were able to observe weak expression of TcKLIF along the FAZ, followed by localization to the new FAZ and the posterior puncta in cells at later cell cycle stages. In cells undergoing cytokinesis, TcKLIF no longer localized to the FAZ but instead was clearly associated with the position of the ingressing furrow. At the latest stages of cell division, TcKLIF localized to two discrete foci at the cell posterior that appeared to be subdomains of the 1B41 punctum (**Fig 8B**). This argues that the 1B41-positive punctum may represent a microtubule-containing structure that replicates during the last stages of cell division, perhaps to facilitate the construction of a new posterior end. The levels of TcPLK1 and TcKLIF signal were not sufficient for U-ExM analysis, which creates a 125-fold decline in fluor density.

Considering the size difference between the two daughter cells produced during *T. cruzi* epimastigote cell division, it is possible that there would be a difference in their cell division rates during their subsequent round of division. We recently developed a strategy using agarose microwells that allows us to confine and image trypanosomatids in small volumes without immobilization, which permits the parasites to divide at rates similar to those in bulk cultures [72]. During optimization of our microwell approach for imaging *T. cruzi* epimastigotes, we found that increasing the height of the wells to 6 microns to account for the width of *T. cruzi* epimastigote cells undergoing cytokinesis provided unimpeded cell divisions, with no loss in viability out to 48 h. We isolated single cells in wells and monitored them until they divided, then tracked the subsequent division rate of the daughter cells (**Fig 9A and S1 Movie**). Cells divided in the microwells at a faster rate than in bulk cultures, with an average rate of 15.4 h for all the events we observed, compared to 22.1 h in culture. This increased rate may reflect the fact that the agarose microwells do not exchange gas with the environment, which would lead to decreased oxygen levels within the well over time. Recent work has shown that *T. cruzi* epimatigotes grow more rapidly in anoxic environments, which is consistent with the conditions within the insect gut that they inhabit throughout their time within this host [73]. We found that there was a 4.9 h difference in the rates of daughter cell divisions in epimastigote *T. cruzi,* with the median first division occurring at 12.8 h and the second division occurring at 17.7 h (**Fig 9B and 9C**). This difference is larger than what we observed in *T. brucei*, which suggests that the increased asymmetry in *T. cruzi* epimastigote cell division leads to a larger difference in daughter cell division rate.

**Fig 9.**
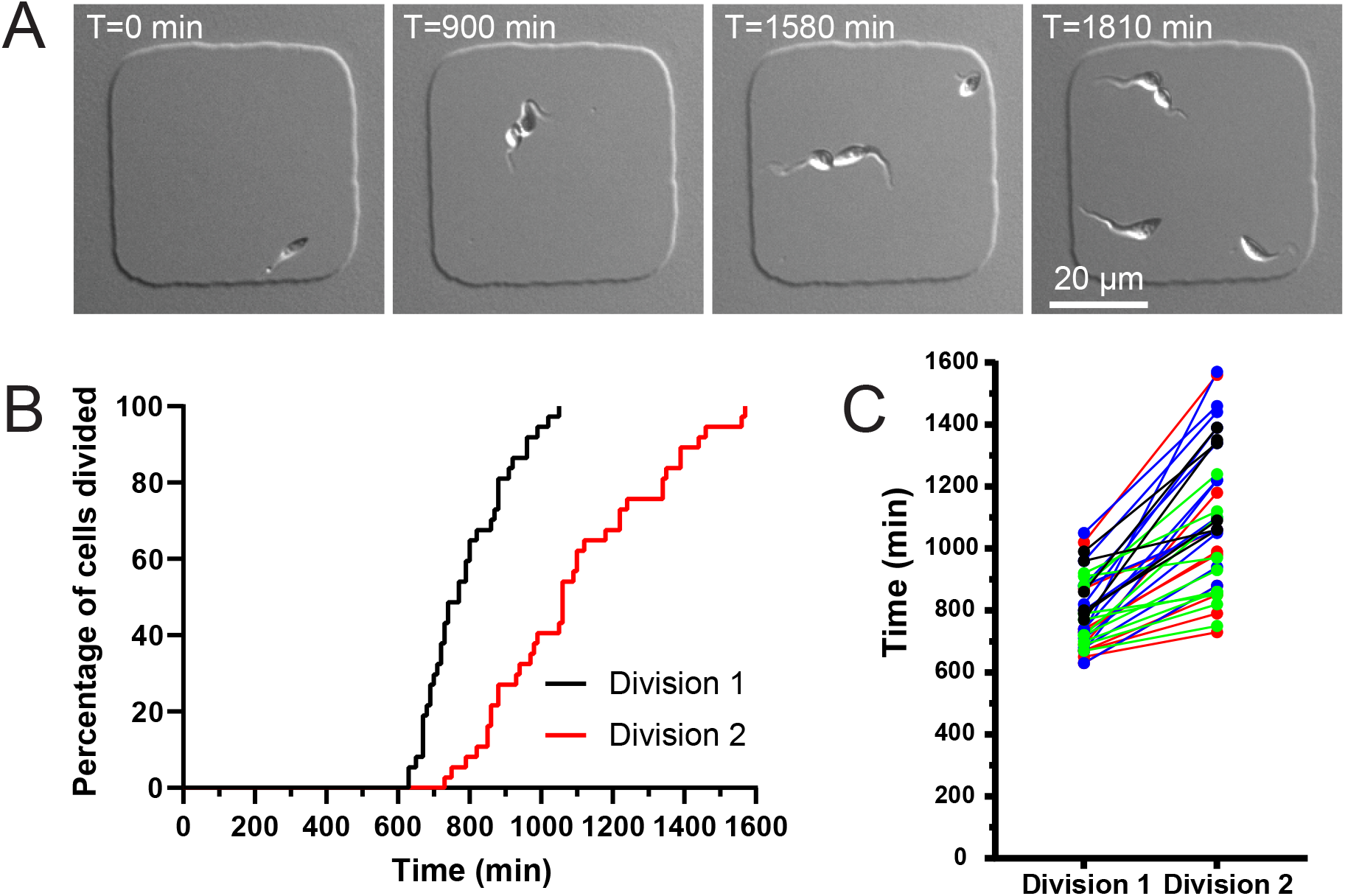
*T. cruzi* daughter cells divide at different rates. (**A**) *T. cruzi* epimastigotes cells were confined in agarose microwells and then imaged using DIC microscopy. Images were taken every 10 min for 2 d. Wells containing one cell were identified and the timing of the first round of division was noted. The time that each daughter cell took to complete a subsequent round of division was recorded. Images show representative still shots of live *T. cruzi* cells undergoing division. In this example, over 220 minutes (3 h, 40 min) elapsed between the division of the first daughter cell and the second. (**B**) Staircase plot showing the timing of the first and second daughter cell divisions. T1/2 for division 1 was 12.8 h, while T_1/2_ for division 2 was 17.7 h. (**C**) Plot showing direct comparison of daughter cell division rates. Each line represents daughter cell division times from the same progenitor cell. Cell comparisons are color coded to depict cells imaged in the same live-cell imaging run.

## Discussion

In this work, we have studied *Trypanosoma cruzi* epimastigote cell division using *T. cruzi* gene editing and components of the parasite cell division apparatus and cytoskeleton that were identified in localization studies conducted in other trypanosomatids. These advances have allowed us to define different *T. cruzi* cell cycle stages based on cell morphology and the localization of marker proteins for comparison with previous work on promastigote *Leishmania* and trypomastigote *T. brucei* stages. Of these four trypanosomatid forms, epimastigote *T. cruzi* appears to have the most asymmetric cell division mechanism, producing a new-flagellum daughter cell only half the length of the old-flagellum daughter cell body. We observed a significant difference in the division rates of the two daughter cells in agarose microwells, which argues that this asymmetry impacts the duration of the subsequent round of cell division. Most of the cytoskeletal and cell-division proteins we studied have altered localization patterns compared to what is observed in *T. brucei* (**Fig 10**). The tip of the growing new FAZ, which recruits many essential cytokinetic proteins such as such as TbTOEFAZ1, TbKLIF, and TbPLK in *T. brucei* does not function as their primary localization point in *T. cruzi.* These proteins appear to localize along the lengths of the old and new FAZ. The posterior end of the cell recruits many of these cytokinetic proteins and may contain tubulin posttranslational modifications that are confined to the FAZ in *T. brucei*. These changes likely reflect the differing geometries that *T. brucei* trypomastigote and *T. cruzi* epimastigote cells adopt during cell division, which put unique constraints on the timing and positioning of duplicating organelles.

**Fig 10.**
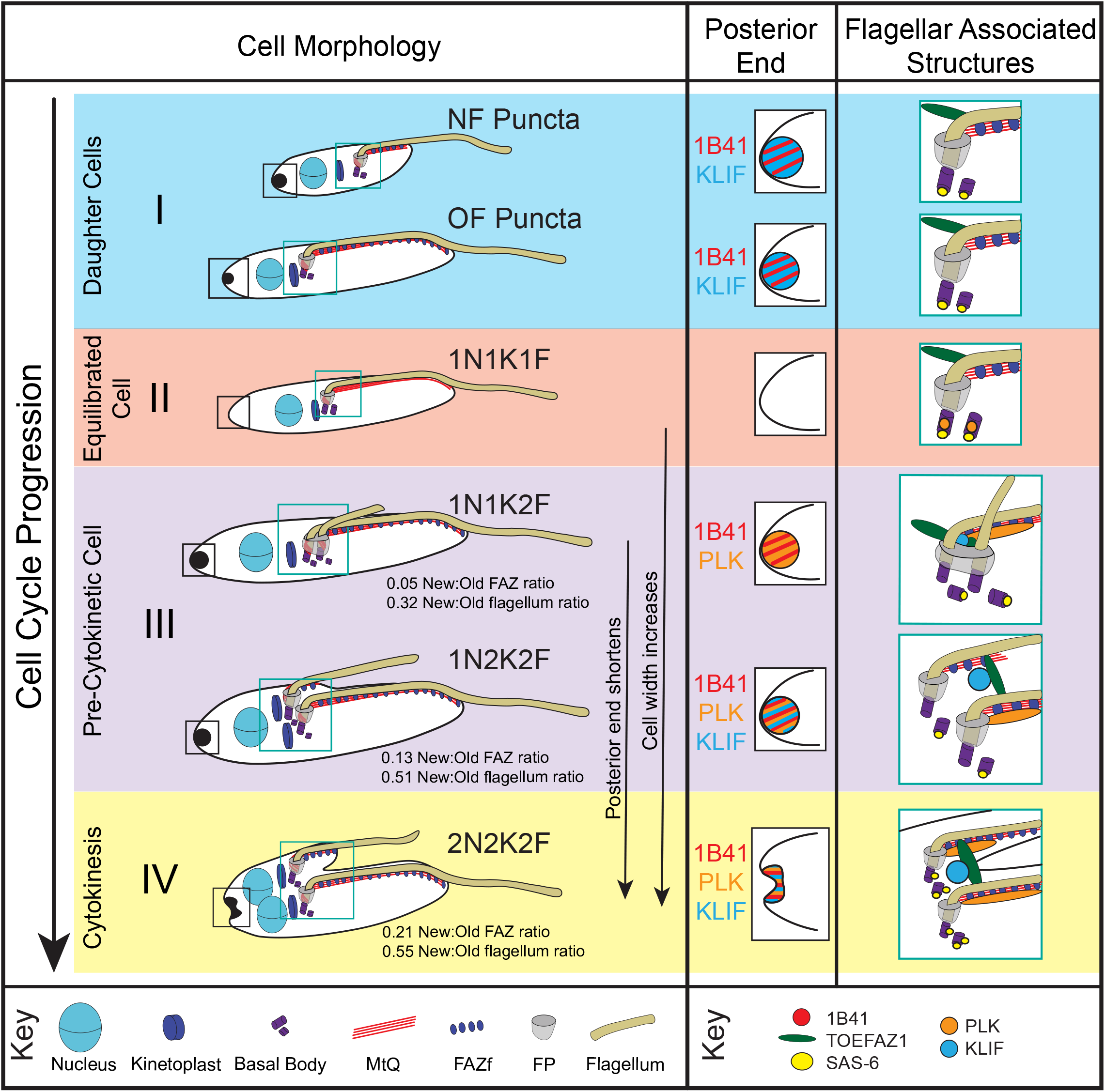
Model describing the different stages of epimastigote *T. cruzi* cell division described in this work. (**I**) At the end of cytokinesis, two types of cells are observed: new flagellum daughters (NF Puncta) that are smaller, have a shorter flagellum and FAZ, and a more intense posterior punctum staining with 1B41 and KLF when compared to the old flagellum daughters (OF Puncta). In both cell types the pro and mature basal bodies are labeled with SAS-6 and TOEFAZ1 is present on the dorsal side of the flagellum. **(II)** Once the new flagellum daughter has extended its FAZ and flagellum, the two daughter cells are indistinguishable. The posterior punctum labeling disappears entirely. As the cell begins to divide, PLK is present on the basal body. TOEFAZ1 still labels the dorsal region. **(III)** As the new flagellum is assembled, both PLK and 1B41 labeling become apparent at the cell posterior. PLK is present along the FAZ as well. SAS-6 labeling appears primarily associated with the probasal bodies. The cell begins to widen along its short axis. As the new flagellum and FAZ continue to grow, the KLIF labeling appears at the site where the cleavage furrow will initiate from and is also present at the posterior punctum. TOEFAZ is recruited to cleavage furrow ingression point. Cell widening continues, while the cell posterior shortens along its long axis. **(IV)** As cytokinesis begins, the posterior punctum is labeled with 1B41, PLK, and KLIF and begins to resolve into two discrete structures. KLIF and TOEFAZ are present at the ingressing cleavage furrow, while PLK is present on both FAZ. SAS-6 labeling reappears at the mature basal body.

Work that predates the development of Cas9 editing in *T. cruzi* examined different aspects of the epimastigote cell cycle, with an emphasis on flagellar pocket and nuclear duplication [74], noted that the new flagellum appeared significantly shorter than the old during the later stages of cell division. One point of difference is that the authors state that the new flagellum length reaches near parity with the old flagellum prior to the completion of cell division. In our experiments, the asymmetry present during the later stages of cell division is carried on to the daughter cells, with the new-flagellum daughter having a significantly shorter new flagellum and FAZ.

Labeling with the 1B41 antibody provides a useful method for tracking division in *T. cruzi*. While the specific epitope of the antibody is still unknown, it appears to be a PTM of beta tubulin. *T. cruzi* cells contain several MT structures that likely derived from the MT rootlets found in the feeding apparatus of the free-living ancestor of the trypanosomatids, including a second MT quartet and a MT triplet associated with the cytopharynx [24,25]. None of these MT structures appear to label with 1B41, suggesting that the FAZ-associated MtQ is uniquely modified. It is currently unknown why the 1B41 signal partitions into the daughter cells asymmetrically, but it may reflect differences in the PTM status of certain microtubule plus ends that selects them for incorporation into the new-flagellum daughter cell posterior, while leaving the less-modified plus ends in the old structure. It is possible that the 1B41 epitope facilitates recruitment of the proteins responsible for duplicating the cell posterior. TcPLK localizes to the cell posterior just prior to the appearance of the 1B41 signal, so the kinase may function as an upstream regulator of this process. KLIF localizes to the cell posterior during late cytokinesis and appears to resolve into two distinct foci, which suggests that the kinesin may be responsible for sorting the MT plus ends into two discrete structures to complete cytokinesis. This is consistent with the proposed function of KLIF in *T. brucei*, where depletion of the motor leads to a block in assembly of a new cell posterior and subsequently prevents cell division [69,70]. In trypomastigotes, formation of a new cell posterior occurs de novo at a location which initially appears as a new focus of MT plus ends that are subsequently gathered into a MT bundle to complete cell division [36]. In epimastigotes, it appears that cell posterior assembly occurs at the site of the old posterior and may involve the addition of MTs, followed by partitioning of the duplicated structure. A previous survey of trypansomatid morphologies has suggested that cells with detached flagella, termed “liberforms”, tend to undergo cell widening during cell division, whereas cells with attached flagella, termed “juxtaforms” lengthen their cells as they divide [7]. It was suggested that epimastigotes, which are juxtaforms, may employ a more trypomastigote-like mechanism for cell division. Our results with *T. cruzi* suggest that epimastigotes actually employ a liberform-like division mechanism. However, limiting the length of the new FAZ, which would create a significant impediment to altering the shape of the cell body, may be necessary to widen the cell. This may explain the extreme asymmetry of cell division in epimastigote *T. cruzi*.

In *T. brucei*, the tip of the new extending FAZ plays an important role in designating the point of cleavage furrow ingression [11]. The tip is in close proximity to the old FAZ and contains a host of proteins that are thought to play a role in triggering cytokinesis, including the CIFs, TOEFAZ1, KLIF, and PLK. In *T. cruzi* epimastigotes, PLK localizes along the old and new FAZ prior to moving to the cell posterior, while KLIF localizes to both FAZ, then the cell posterior, followed by the cleavage furrow. While both of these proteins remain cell-cycle regulated, TOEFAZ1 appears to be persistently expressed, initially localizing to a position similar to where the new FAZ is assembled, then moving along the ingressing furrow. In trypomastigote *T. brucei*, the extension of the new flagellum and FAZ are tightly coordinated, with the FAZ extending 1-2 microns behind the flagellum [75]. In *T. cruzi* epimastigotes, FAZ and flagellar growth are not tightly coupled, as the new flagellum extends far beyond the end of the new FAZ. It is possible that using the tip of the new FAZ to target the cytokinetic complex may be a feature exclusive to cells with trypomastigote morphology.

The basal body protein TcSAS-6 appears to have a different localization pattern in *T. cruzi* compared to other trypanosomatids. The protein is one of the earliest components of the probasal body and is thought to establish the 9-fold symmetry of the organelle by constructing a cartwheel-shaped structure at its base [55,76,77]. In *T. brucei*, depletion of TbSAS-6 causes defects in probasal body assembly and blocks the production of a new flagellum, causing growth arrest [56]. In some organisms, the cartwheel structure breaks down during the completion of cell division, which leads to a loss in SAS-6 signal, although the cartwheel remains as a persistent component of the basal body in many organisms including *T. brucei* [78]. In *T. cruzi*, TcSAS-6 is present at the pro and mature basal body early in the cell cycle, then moves to the bases of the assembling probasal bodies. It then is restored to the mature basal bodies prior to cytokinesis while remaining associated with the probasal bodies. This inheritance pattern is consistent with the function of SAS-6 in probasal body assembly and may suggest that the new cartwheel is assembled on the basal and probasal bodies after their duplication but prior to the completion of cell division, so that the new probasal bodies can be assembled during the subsequent round of division. While this would be different than what is observed in *T. brucei*, earlier reports have shown that TbSAS-6 levels on the basal body fluctuate during cell division, with higher levels in the probasal body during its assembly [79].

Our localization and morphology studies have highlighted significant differences in *T. cruzi* epimastigote cell division compared to previously studied trypanosomatids. The next step will be to determine if the changes in localization of proteins such as TcPLK and TcKLIF reflect changes in their function. While knockout strategies are available in *T. cruzi*, methods for targeting essential genes using conditional or inducible approaches are still under development. It would also be interesting to localize these proteins in *T. brucei* epimastigotes to determine if they retain the localization pattern seen in trypomastigotes or if their morphotype alters the patterns. Currently, *T. brucei* epimastigotes can only be isolated directly from the insect host [31,33]. As the ancestral trypanosomatid likely had a promastigote-like morphology, its cell division mechanism served as a starting point for diversification as organisms moved into new hosts [80]. The trypomastigote form appears to be primarily an adaptation to survival within the bloodstream of vertebrate hosts, which has made it a focus of much of the previous work on trypanosomatids. However, with a better understanding of the range of morphologies present among these organisms and the tools for sequencing and manipulating them, a broader understanding of their cell division pathways is now feasible.

## Abbreviations

FAZ: flagellum attachment zone
PTM: posttranslational modification
MT: microtubule
U-ExM: Ultrastructure expansion microscopy

## Acknowledgements

We would like to thank Rick Tarleton for providing *T. cruzi* cells, Martin Taylor for the Cas9-pLEW13 vector, Jack Sunter for the TAT1 antibody, and Kate O’Connor Giles and the Carney Institute of Brain Science for access to the Nikon SoRa microscope. We would also like to thank Sue Vaughan and Drew Etheridge for valuable insights. The research reported in this publication was supported by NIAID-NIH under awards R01AI166363 and R21AI151490 to CLdG. The content is solely the responsibility of the authors and does not necessarily represent the official views of the National Institutes of Health.

## Materials and methods

### Cell culture and growth curves

Cells used for all experiments were *T. cruzi* Y strain epimastigotes were cultured in LDNT medium (0.5% Oxoid neutralized liver digest (ThermoFisher Scientific, Waltham, MA), 68 mM sodium chloride (ThermoFisher Scientific), 70 mM Tryptone (ThermoFisher Scientific), 5 mM potassium chloride (ThermoFisher Scientific), 58 mM sodium phosphate dibasic (ThermoFisher Scientific), 11 mM glucose (ThermoFisher Scientific), 25 μM Hemin (Sigma-Aldrich, St. Louis, MO), gentamicin (25 μg/mL) (ThermoFisher Scientific), 10% heat inactivated fetal bovine serum (Bio-Techne), penicillin-streptomycin (100U/mL) (ThermoFisher Scientific), pH 7.2) at 27 °C. For selecting cell lines, blasticidin was used at 100 µg/mL and G418 at 500 µg/mL. Cell counts were obtained with a Z2 Coulter Counter (Beckman Coulter, Brea, CA). Growth curves were performed in triplicate seeding a culture of cells at 2×10^6^ cells/mL. Cell counts were obtained every 24 h for 9 d with reseeding 2×10^6^ cells/mL into fresh flasks and media every 3 d.

### Calculation of cell cycle timing

In order to calculate relative time spent in each cell cycle intermediate, a method previously described [37,54]was employed to account for an asynchronous logarithmic culture growth of cells that divide by binary fission with the equation:

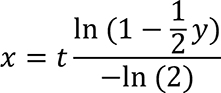

In this equation, x= the time through the cell cycle, y= proportion of cells preceding and including the cell cycle stage in question, and t= cell cycle length.

### Cloning and cell line assembly

Cas9 and T7 RNA polymerase expressing cells were generating using a pLEW13-Cas9 plasmid provided by Martin Taylor (London School of Hygiene and Tropical Medicine, London, United Kingdom) as previously described [17]. Healing constructs for gene tagging were generated by PCR using gene-specific primers with 30 bp homology flanking the CRISPR-Cas9 cut site. A base plasmid containing a blasticidin resistance marker, an intergenic sequence isolated from *T. brucei*, and a 3×Ty1 epitope tag fused to a mNeonGreen fluorescent protein was employed as a template for PCR to generate all healing constructs All proteins studied here were tagged as closely to the N-terminus as allowed by available CRISPR-Cas9 cut sites. PCRs to generate the healing constructs were performed in triplicate, pooled and purified with a Zymo Research DNA Clean and Concentrator kit (Genesee Scientific, San Diego, CA) and resuspended in 25 μL of ultra-pure water (Genesee).

sgRNAs were prepared with a gene specific forward primer with target site determined by Eukaryotic Pathogen CRISPR guide RNA/DNA Design tool (http://grna.ctegd.uga.edu/) using the *T. cruzi* Y C6 Pacbio genome assembly [14,15]. sgRNA was amplified in triplicate as previously described[81]. sgRNA PCRs were pooled and purified with a Zymo Research DNA Clean and Concentrator kit (Genesee Scientific) and resuspended in 25 μL of Ultra-Pure Water (Genesee). Purified sgRNA was pooled with corresponding purified healing template for a total of 50 μL. Primers used in this study can be found in S1 Table.

5×10^7^ cells expressing SpCas9 and T7 RNA polymerase from a logarithmically growing culture were harvested by centrifugation at 800×g for 5 min at 4 °C. Cells were resuspended in 225 μL nucleofection buffer (90 mM sodium phosphate dibasic, 5 mM potassium chloride, 0.15 mM calcium chloride hexahydrate, 50 mM HEPES). Pooled sgRNA and healing template were added to the cell suspension and nucleofected using a Lonza Nucleofector 2b (Lonza, Basel, Switzerland) using program X-014. Cells were added to 10 mL LDNT media without selection markers. 24 h post nucleofection, selection marker was added to the cells. Once a polyclonal bulk culture was established, cells were plated for clonality by serial dilution.

### Immunofluorescence

Cells were fixed in media with an equivalent volume of 8% PFA in PBS (4% final concentration) for 20 min at RT. Cells were then centrifuged at 800×g for 5 min at RT and washed once in 1 mL of PBS. Cells were centrifuged onto coverslips at 800×g for 5 min 4 °C. Coverslips were then inverted onto a solution of 0.5% NP-40 in PBS for 5 min, followed by 3×5 min washes in PBS. Cells were incubated in blocking buffer (5% goat serum (Gibco, ThermoFisher Scientific, Waltham, MA) in PBS) followed by 3×5 min washes in PBS. Coverslips were incubated in primary antibody in blocking buffer for 1 h at RT followed by 3×5 min washes in PBS. Coverslips were then incubated in secondary antibody in blocking buffer for 1 h at RT followed by 3×5 min washes in PBS and then mounted in Fluoromount G with DAPI (Southern Biotech, Birmingham, AL).

### Ultrastructure expansion sample preparation

1×10^6^ cells per gel were fixed in solution by adding 2X fixative solution (1.4% PFA, 2% Acrylamide, 2X PBS) 1:1 to cells in media. Cells were spun 1000×g for 10 min at RT. Cells were resuspended in 750 μL 1X fixative solution (0.7% PFA, 1% acrylamide, 1X PBS) and spun again 1000×g for 10 min at RT. Cells were resuspended in 500 μL of 1X fixative solution and spun down onto 12 mm coverslips by briefly centrifuging at 800×g at 4 °C. Coverslips were inverted onto 100 μL of 1X fixative solution in a humidified chamber and incubated at 37 °C for 3.5 h. Slips were inverted onto 40 μL of gelation solution (19% sodium acrylate, 10% acrylamide, 2% Bis, 1×PBS). 40 μL of fresh gelation solution was added to 0.5 mL centrifuge tubes. 2 μL of 10% tetramethylethylendiamine (TEMED), followed by 2 μL of ammonium persulfate (APS) were added to the 40 μL gelation solution to initiate polymerization. The tube was vortexed briefly, then the solution was pipetted onto 18 mm coverslips. Cells on 12 mm coverslips were then inverted onto the activated gelation solution. Slips were placed at 37 °C to fully polymerize for 1 h in a 12 well plate. 1.5 mL denaturation buffer (200 mM SDS, 200 mM NaCl, 50 mM Tris) was added to the wells and placed on a rocker at RT for 15 min. Gels were removed from cover slips and placed in a 1.5 mL tube with 1 mL denaturation buffer. Gels were denatured at 95 °C for 30 min. Gels were poured into 150 mm petri dishes filled with DI water and allowed to expand for 30 min at RT. Water was exchanged for fresh DI water and allowed to expand overnight. The next day, gels were shrunk in 1X PBS for 30 min. Gels were transferred to a 6 well plate and 300 μL of primary antibody in blocking buffer (2% BSA in 1X PBS) was added and allowed to incubate on a rocker for 8 h. Gels were washed 3×10 min in PBS-Tween on a rocker. 300 μL of secondary antibody in blocking buffer was added and allowed to incubate at 37 °C with gentle agitation overnight. Gels were washed 3×10 min in PBS-Tween on rocker. Gels were then re-expanded in DI water with 2 water changes every 30 min. Next day, a 10 mm diameter gel slice was excised and placed in a MatTek 35 mm dish, No. 1.5 Coverslip, 10 mm Glass Diameter (MatTek Corporation, Ashland, MA) coated with Poly-D-Lysine (Thermo Fisher Scientific) for imaging.

### Antibodies

The following primary antibodies and dilutions were used in these experiments: anti-Ty1 (1:1,000 for IF and western blotting, 1:10 for U-ExM) from Sebastian Lourido (Massachusetts Institute of Technology, Boston, MA), 1B41 (1:1,000 for IF) from Linda Kohl (Centre National de la Recherche Scientifique, Paris, France), TAT1 (1:10,000 for IF, 1:100 for U-ExM) from Jack Sunter (Oxford Brookes University, Oxford, United Kingdom), anti-polyglutamylation GT335 (1:25,000 for IF) (Adipogen, San Diego, CA), anti-polyglutamate chain IN105 (1:10,000 for IF) (Adipogen), 20H5 (1:1000 for IF) from EMD Millipore.

Secondary antibodies were used in these experiments. All dilutions for IF were 1:1000, U-ExM were 1:100, and western blotting were 1:10,000. Goat anti-mouse IgG1 Alexa Fluor 488 (Thermo Scientific), Goat anti-mouse IgM Alexa Fluor 488 (Thermo Scientific), Goat anti-mouse IgG2a Alexa Fluor 568 (Thermo Scientific), Goat anti-rat IgG H&L Alexa Fluor 568 (Thermo Scientific), Goat anti-rabbit IgG Alexa Fluor 568, Goat anti-mouse IgG (H&L) HRP (Thermo Scientific).

### Immunofluorescence microscopy

IF images were obtained on a Zeiss Axio Observer Z1 microscope (Carl Zeiss Microscopy, Oberkochen, Germany) using a 100×/1.4 NA Plan Aprochromat objective lens. Images were captured using an ORCA-Flash 4.0 V2 sCMOS camara (Hamamatsu, Shizouka, Japan) with Slidebook 6 microscopy software (Intelligent Imaging Innovations Inc.)

### SoRa microscopy

Ultrastructure expansion gels were imaged on a Nikon Eclipse Ti2 microscope (Nikon Instruments, Melville, NY) equipped with a Yokogawa CSU-W1 SoRa super resolution confocal scanning unit (CSU-W1, Yokogawa Electric, Musashino, Japan) and a Photometrics Prime BSI sCMOS camera (Teledyne Photometrics, Tucson, AZ) using a CFI APO TIRF 60× 1.49 oil objective (Nikon Instruments). Images were taken as a z-stack at 0.1 μm per step. Images were denoised using Nikon NIS-elements software and deconvoluted using Richardson-Lucy deconvolution.

### Generating agarose microwells

PDMS stamps and agarose microwells were fabricated as previously described, for a final concentration of 3.5% SeaPlaque agarose (Lonza, Basel, Switzerland) in LDNT media [72]. Microwell dimensions used in these studies were 50×50×6 μm. Prepared agarose grids were stored at 4 °C for up to one week with sufficient LDNT media to prevent drying out.

### Live cell plating and imaging

1 mL of *T. cruzi* cells expressing Cas9 and the T7 polymerase at 1×10^7^ cells/mL density were centrifuged at 800×g for 5 min at RT. Cells were resuspended in 1 mL fresh LDNT media with 100 μM reduced L-glutathione (Sigma-Aldrich, St. Louis, MO). 125 μL of cells containing 1.25×10^6^ cells were plated in a Lab-Tek II chambered coverglass #1.5 Borosilicate chamber (Nunc International, Rochester, NY). A cut-to-fit agarose grid was inverted onto the cells and the chamber sealed as previously described. Imaging chamber was placed into a live-cell imaging insert (Oko labs) set to 27 °C and mounted on a Zeiss Axio Observer Z1 microscope using a 20×/0.8 NA Plan Aprochromat objective lens. Cells were imaged every 10 minutes for 48 h with 30 ms exposures. In order to compensate for focus drift over long-term imaging, a Definite Focus 2 system (Carl Zeiss Microscopy) was employed. Images were captured using an ORCA-Flash 4.0 V2 sCMOS camara with Slidebook 6 microscopy software.

### Microscopy analysis

All microscopy images were analyzed with ImageJ (National Institutes of Health, Bethesda, MD). Figures for publication were prepared in Adobe Photoshop and Illustrator (Creative Cloud 2022).

### Statistical analysis

Statistical analysis and graphs were generated using GraphPad Prism 9 software version 9.1.1 (GraphPad Software). For morphometric analysis, one-way ANOVA analysis was performed to identify significant differences (p<0.05) between cell cycle intermediates for each morphometric measurement. For live-cell analysis, a Power analysis was performed to ensure sufficient n was performed to ensure conclusion was reasonable. For live-cell differential daughter division times, an unpaired two-tailed students t-test was performed to determine if differences observed was statistically significant at the p<0.05 significance level. All graphical data was plotted using SuperPlots to show variability among biological replicates [82].

## Supplemental Figure legends

**S1 Fig.**
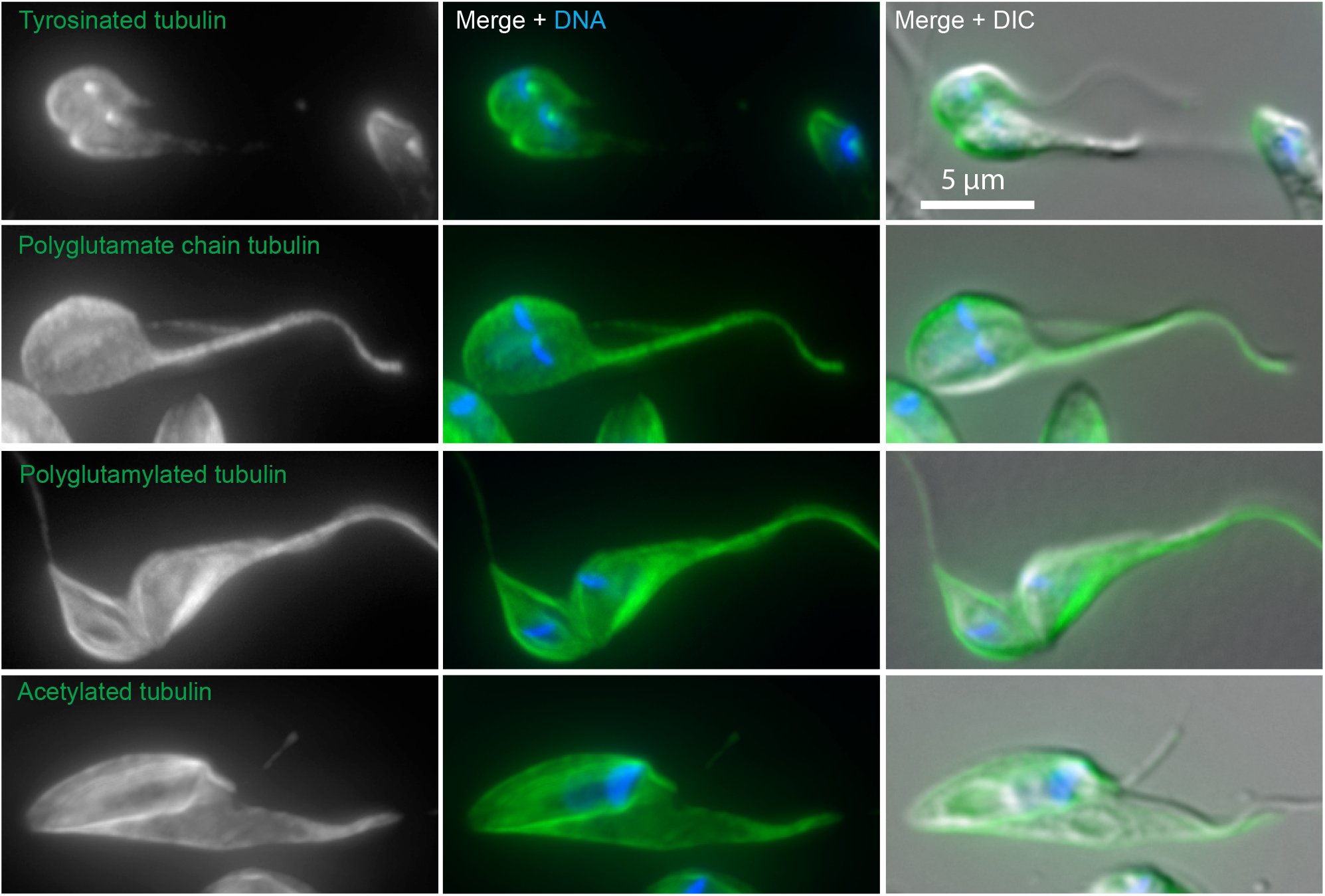
Antibodies against established tubulin post-translational modifications do not label the 1B41-positive posterior punctum. *T. cruzi* cells were fixed, then labeled with anti-tyrosinated tubulin (YL1/2), polyglutamylated tubulin (GT335 or IN105), or acetylated tubulin (6-11 B-1), followed by secondary antibodies conjugated to Alexa488, and DAPI to label DNA. Cells were then imaged using epifluorescence and DIC microscopy.

**S2 Fig.**
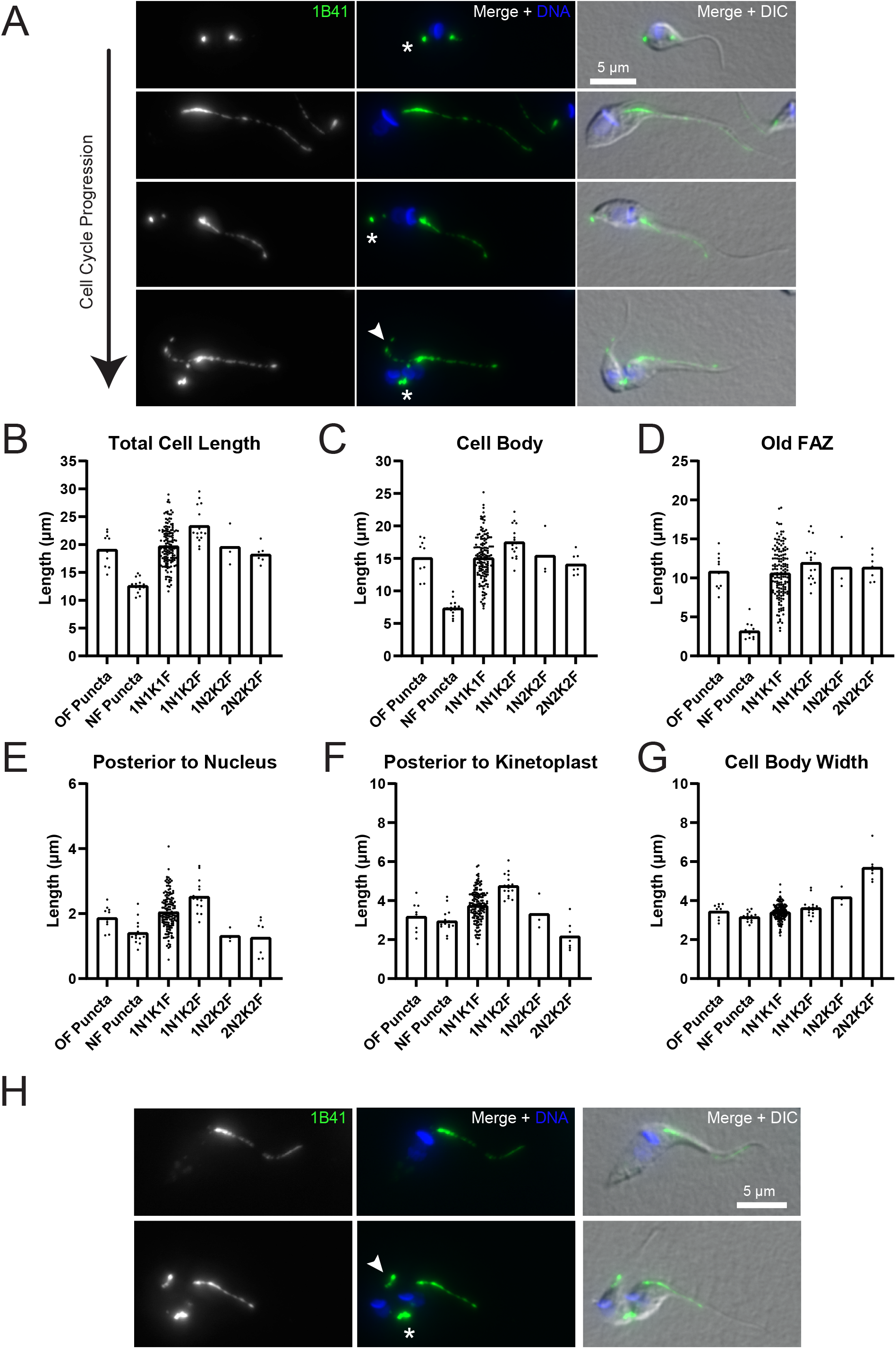
The parental *T. cruzi* Y strain cell line and the *T. cruzi* Brazil A4 strain both show asymmetric cell division and a 1B41-positive punctum at the cell posterior. (**A**) The parental *T. cruzi* Y strain used for making all the tagged cell lines in this work was fixed and then labeled with 1B41 antibody, followed by fluorescently conjugated secondary antibodies and DAPI to label the DNA. The cells were imaged using immunofluorescence and DIC microscopy. Asterisks highlight the 1B41 posterior punctum labeling, while the arrowheads identify the new FAZ. (**B**) Measurements of the total cell length as defined in Fig 4A. (**C**) Measurements of the cell body length as defined in Fig 4A (**D**) Measurements of the Old FAZ length as defined in Fig 4A (**E**) Measurements of the posterior to nucleus distance as defined in Fig 4A. (**F**) Measurements of the posterior to kinetoplast distance as defined in Fig 4A (**G**) Measurements of the cell body with as defined in Fig 4A. (**H**) *T. cruzi* Brazil A4 strain cells were fixed and labeled with 1B41, followed by fluorescently conjugated secondary antibodies and DAPI to label the DNA. The cells were then imaged using immunofluorescence microscopy.

**S3 Fig.**
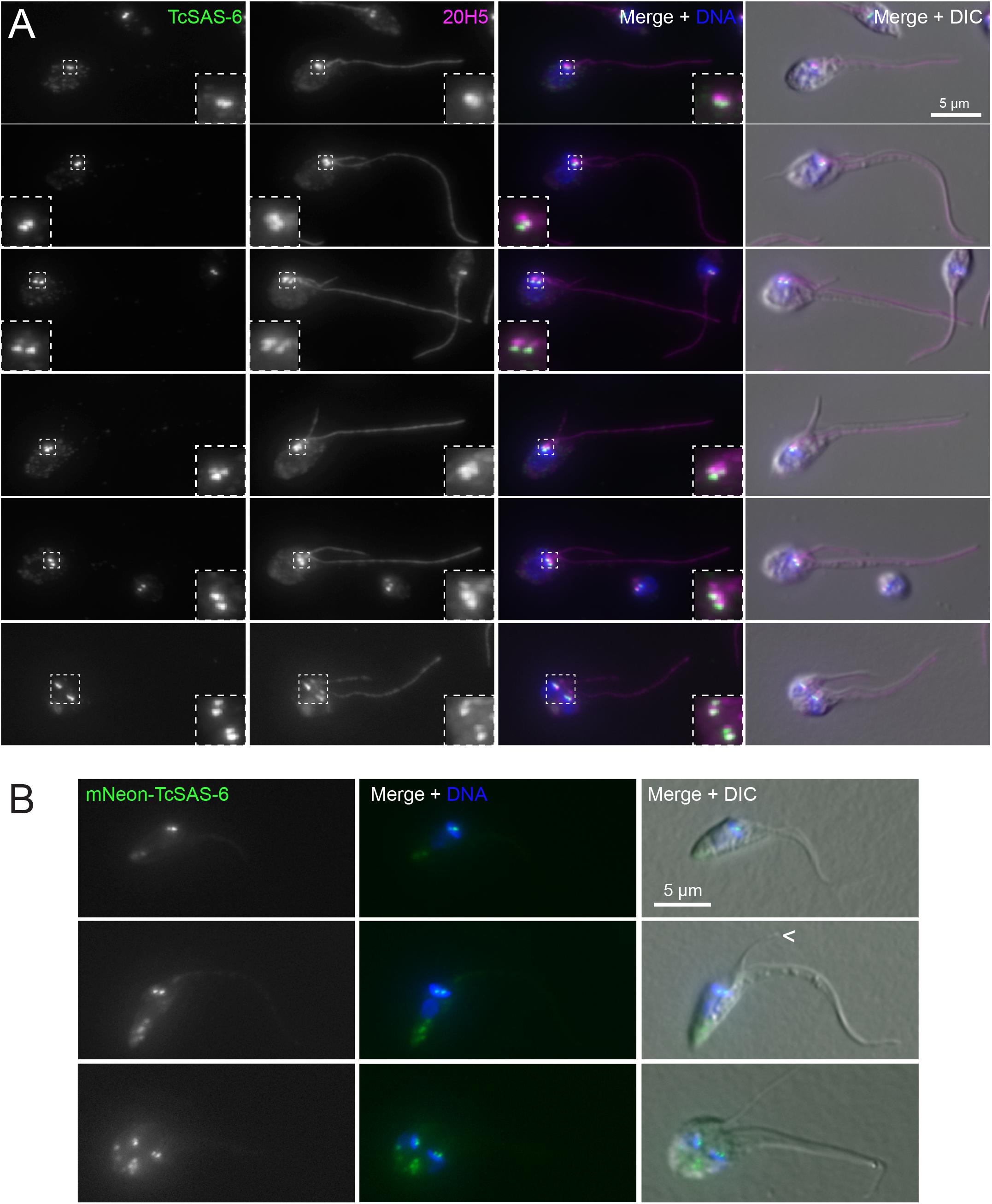
TcSAS-6 does not persistently label the pro- and mature-basal body in *T. cruzi* epimastiogtes. (**A**) Cells carrying a 3xTy1-mNeonGreen::TcSAS-6 allele were fixed and labeled with anti-Ty1 and 20H5 to label SAS-6 and TcCentrin2, followed by fluorescently labelled secondary antibodies and DAPI to label DNA. The cells were then imaged using immunofluorescence microscopy. Insets show a 3X magnification of the basal body regions. (**B**) Cells carrying a 3xTy1-mNeonGreen::TcSAS-6 allele were fixed and then incubated with DAPI to label the DNA. The innate mNeonGreen fluorescence was then imaged using epifluorescence microscopy.

**S4 Fig.**
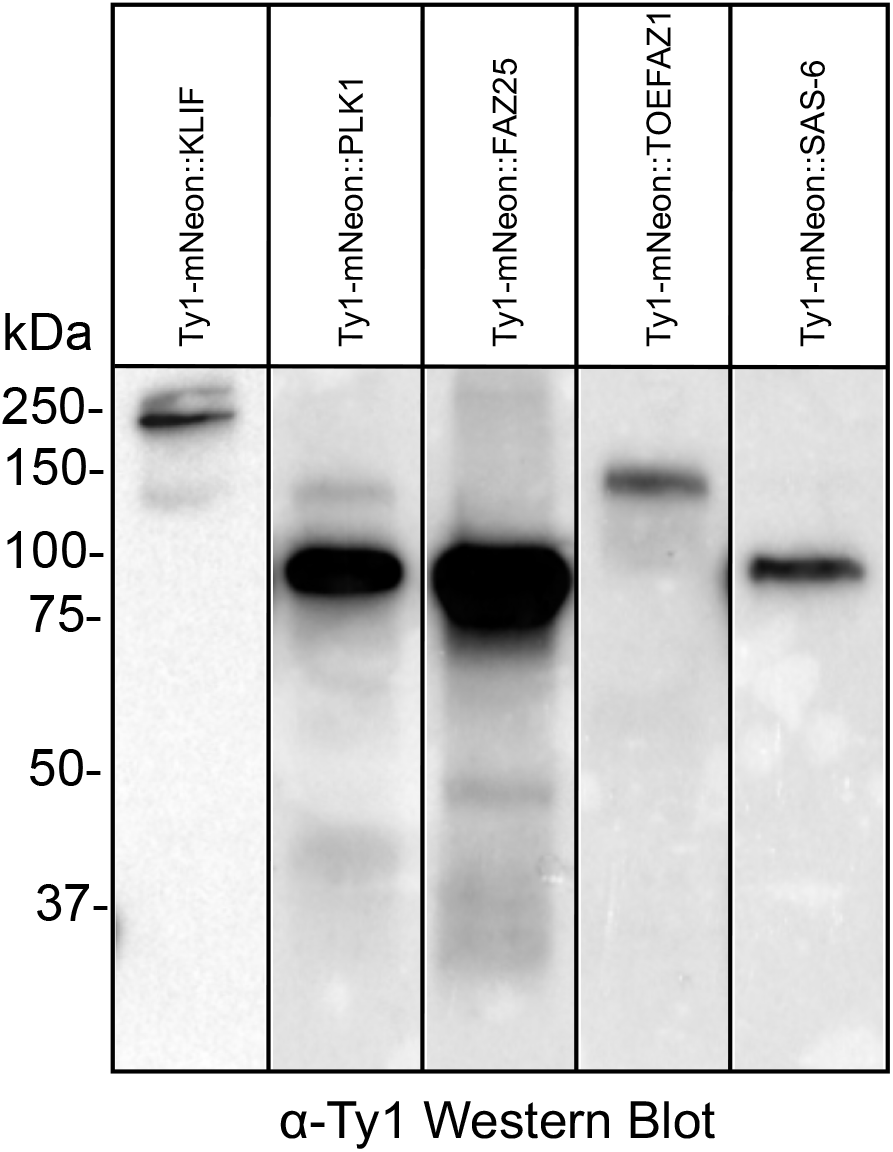
Western blotting validation of the tagged cell lines used in this work. The KLIF, PLK, FAZ25, TOEFAZ1, and SAS-6 cell lines endogenously tagged with 3X Ty1-mNeonGreen were harvested and then lysed in SDS-PAGE loading buffer, followed by fractionation using SDS-PAGE. Fractionated lysates were transferred to nitrocellulose and probed with anti-Ty1 antibody.

**S1 Movie. *T. cruzi* epimastigote cell division produces daughter cells with very different rates of subsequent division events.** *T. cruzi* Y strain cells expressing Cas9 and T7 polymerase were plated in 50×50×6 μm agarose microwells and imaged with a 20×/0.8 NA lens. DIC images were captured every 10 min for 48 h. Video depicts images portrayed in Fig 9A.

**S1. Table. All the primers that were used to generate gDNAs and the healing constructs for gene tagging.**

